# Small tau aggregates exhibit disease-specific molecular profiles across tauopathies

**DOI:** 10.1101/2025.06.10.658934

**Authors:** Dorothea Böken, Melissa Huang, Yunzhao Wu, Emre Fertan, Jeff Y.L. Lam, Georg Meisl, Shaan Baig, James B. Rowe, Colin Smith, Annelies Quaegebeur, Dezerae Cox, William A. McEwan, David Klenerman

**Affiliations:** Yusuf Hamied Department of Chemistry, University of Cambridge, Cambridge, CB2 1EW UK; UK Dementia Research Institute, University of Cambridge, Cambridge, CB2 0AH UK; Division of Life Science and State Key Laboratory of Molecular Neuroscience, The Hong Kong University of Science and Technology, Hong Kong; Department of Clinical Neurosciences and Cambridge University Hospitals NHS Trust, University of Cambridge, Cambridge, CB2 0SZ, UK; Department of Neuropathology, University of Edinburgh, Edinburgh, EH4 2XU, UK; School of Sciences, University of Wollongong, Wollongong, NSW 2522, Australia

## Abstract

Tauopathies are neurodegenerative diseases marked by pathological tau aggregation. While disease-specific folds of insoluble tau filaments have been established, it remains unclear whether the smaller, earlier species also differ across tauopathies. Here, we characterise these small tau aggregates from post-mortem brain of individuals with Alzheimer’s disease (AD), progressive supranuclear palsy (PSP), corticobasal degeneration, Pick’s disease, and healthy controls. Using two complementary single-molecule assays, we confirm that small tau aggregates vary in abundance, morphology, and post-translational modifications. AD features specific long, fibrillar-shaped aggregates enriched in phospho-epitopes, while PSP aggregates are shorter, round, and selectively phosphorylated at serine-356, a site we identify as correlating with markers of inflammation and apoptosis. Aggregate properties co-vary with cellular stress signatures and align with disease-specific seeding profiles, suggesting distinct pathological mechanisms. These findings suggest that small tau aggregates are not a shared intermediate, but instead encode disease-specific mechanisms, with potential as both biomarkers and therapeutic targets.

**Key points:** Small tau aggregates show disease-specific signatures across tauopathies, differing in abundance, morphology, and post-translational modifications.

Tau aggregates in AD show enhanced phosphorylation density and structural heterogeneity, including a distinct population of long fibrillar species detectable in the soluble fraction.

Alzheimer’s disease is characterised by specific long, fibrillar-shaped tau aggregates enriched in disease-relevant phospho-epitopes.

PSP features round pSer356-positive aggregates that correlate with apoptotic and inflammatory markers, suggesting a distinct mechanism of toxicity.

Isoform-specific biosensor assays reveal divergent seeding behaviour: CBD shows strong 4R seeding, while PSP lacks seeding activity.

Features of small aggregates co-vary with distinct patterns of gliosis and cell stress, suggesting disease-specific mechanisms of tau-mediated toxicity.

## INTRODUCTION

Tauopathies are a group of neurodegenerative disorders characterised by the pathological aggregation of the microtubule-associated protein tau in the brain^1^. These include Alzheimer’s disease (AD), progressive supranuclear palsy (PSP), corticobasal degeneration (CBD) and Pick’s disease (PiD). Under physiological conditions, tau dynamically associates with microtubules, modulates their stability, and supports axonal transport^2,3^. In disease, tau becomes aberrantly phosphorylated and aggregates, losing its normal function and gaining toxic properties that contribute to neuronal dysfunction and death^4^.

Despite pathological tau aggregation being a shared hallmark of tauopathies, the diseases are pathologically distinct^5^. AD, the most prevalent tauopathy, features a mixture of 3-repeat (3R) and 4-repeat (4R) tau pathology and is considered a secondary tauopathy as tau aggregation is facilitated or accelerated by prior beta-amyloid (Aβ) pathology^6,7^. Clinically, AD is primarily characterised by cognitive decline, with early impairment in declarative and working memory. In contrast, PSP and CBD are primary tauopathies that are typically associated with shorter disease duration compared to AD^8,9^. Both give rise to mixed movement disorders, with a variable degree of cognitive impairment that may precede or follow motor deficits. PSP and CBD are characterised by prominent glial tau pathology in astrocytes and oligodendrocytes, in addition to neuronal tau inclusions^10,11^. PSP has tufted astrocytes, while CBD has astrocytic plaques. In contrast, PiD, a rare form of frontotemporal dementia, is defined by 3R tau pathology and the presence of intraneuronal Pick bodies. Tau inclusions have also been reported in astrocytes in PiD, though these are less prominent than in PSP and CBD and their role remains poorly understood^12^. While reactive astrocytosis is observed in PiD, it may be a response to neuronal loss^13,14^, in contrast to PSP and CBD. PiD often manifests in midlife and progresses rapidly^15^. In addition to differences in tau isoform composition and astroglial involvement, emerging evidence suggests that disease-specific patterns of neuroinflammation may also contribute to divergent pathogenic trajectories across tauopathies^16,17^. The factors underlying these disease-specific differences remain unclear but may involve differences in tau isoform expression, post-translational modifications (PTMs), local cellular environments, or co-pathologies^5^.

Recent advances in cryo-electron microscopy (cryo-EM) have revealed that each tauopathy is characterised by distinct tau filament structures, with highly conserved disease-specific folds of the insoluble aggregates^18–20^ (Schematic 1A), suggesting that conformational differences underlie their divergent clinical and pathological phenotypes^9^. While these larger, insoluble inclusions have historically been the focus of tauopathy research, growing evidence suggests that small, diffusible tau aggregates are the more toxic forms^21^ that play a central role in disease development^22–24^. For example, these small aggregates can impair synaptic function^25,26^, disrupt protein homeostasis^27^ and trigger neuroinflammation^28^. In several recent models, diffusible aggregates correlate more strongly with neurodegeneration than insoluble tau tangles^29^; in contrast, the insoluble species may potentially serve as protective ‘sinks’ that sequester the freely diffusing oligomers^30^, though they may also contribute to cytotoxicity. However, the low abundance, structural heterogeneity, and lack of specific molecular markers of these oligomers have made them exceptionally difficult to study^31^. As a result, it remains unclear whether these small aggregates exhibit disease-specific differences, or whether they represent a common cytotoxic intermediate across tauopathies. In this study, we refer to these smaller “soluble” tau aggregate species as “small tau aggregates” to distinguish them from insoluble fibrils. They are retained in the low-detergent fraction at medium speed centrifugation, exhibit multivalency, and are variously described in the literature as “soluble,” “diffusible,” or “oligomeric” tau. The significance of the variation in size, within and between tauopathies, is uncertain.

**Schematic 1.**
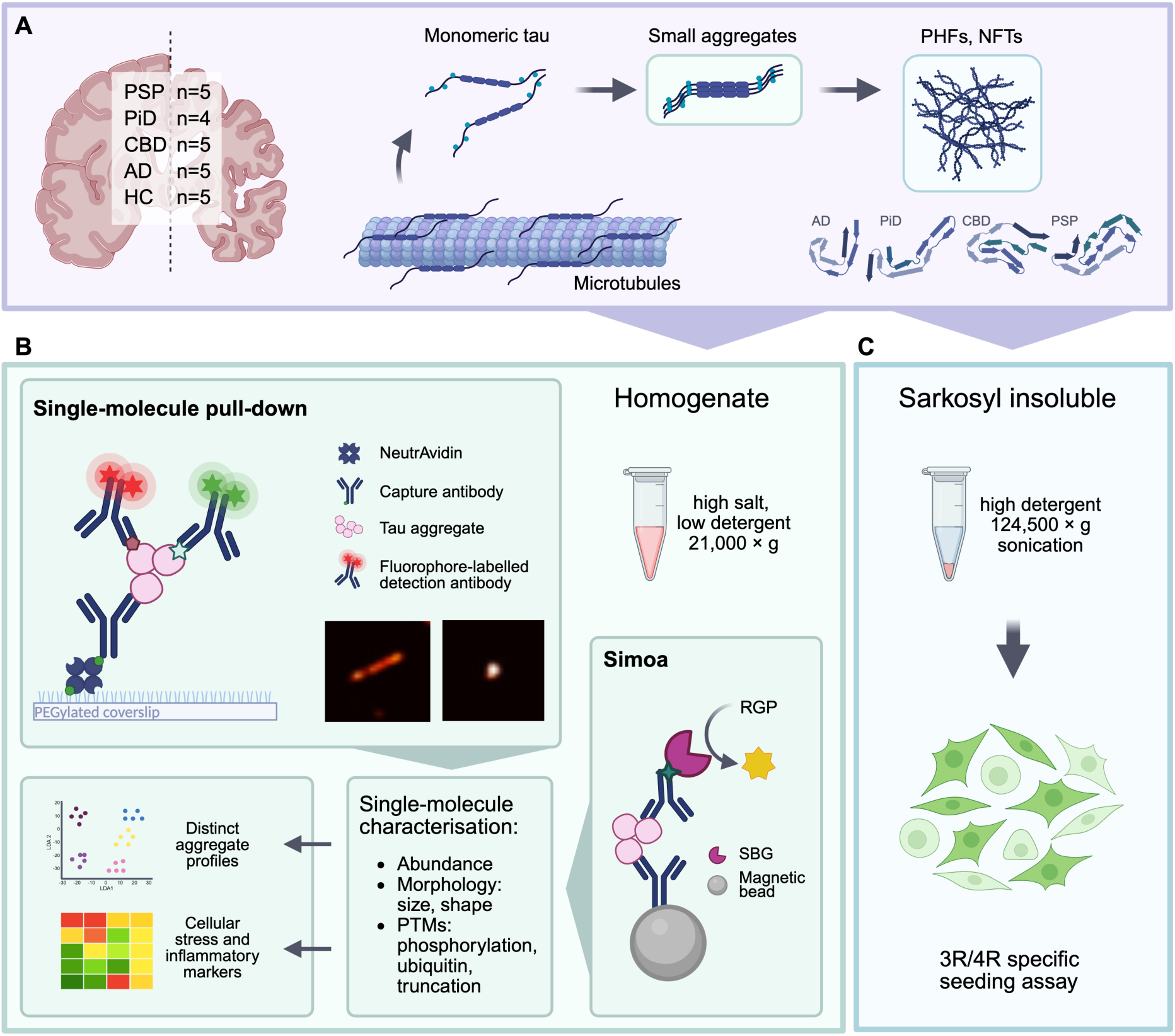
: **Schematic overview of experimental strategy for tau aggregate characterisation across tauopathies.** (A) Tau aggregation proceeds from monomeric tau through small aggregates to mature, fibrillar inclusions such as paired helical filaments (PHFs) and neurofibrillary tangles (NFTs). Distinct folds of the late-stage insoluble tau filaments are associated with different tauopathies, as revealed by cryo-EM. In this study, we asked whether disease-specific differences are already apparent at the level of small tau aggregates. To address this, we analysed tau species retained in the brain homogenate (high salt, low detergent). (B) Small tau aggregates were profiled using two complementary single-molecule methods. The single-molecule pull-down assay captures and images individual tau aggregates using total internal reflection fluorescence (TIRF) microscopy, enabling direct visualisation and quantification of their morphology and post-translational modifications. In parallel, Simoa provides digital, ultrasensitive quantification of epitope-specific aggregates using bead-based immunoassays. Both approaches use a monoclonal antibody sandwich that selectively detects tau species containing at least two epitopes (≥ dimers), excluding monomeric tau. (C) Sarkosyl-insoluble aggregates were assessed in a 3R/4R tau biosensor assay to measure isoform-specific seeding capacity. Aggregate features were linked to disease-specific profiles of cellular stress and inflammatory markers, enabling mechanistic insights into tauopathy heterogeneity.

To address this gap, we systematically characterised the small tau aggregates formed during the earlier stages of aggregation extracted from post-mortem brain of individuals with AD, PSP, CBD, PiD, and age-matched controls without any disease pathology, all sampled from the same brain region (frontal cortex). We applied two complementary single-molecule methods: MAPTau, a tau aggregate pull-down assay coupled with total internal reflection fluorescence (TIRF) microscopy^32^, and tau aggregate-specific single-molecule array (Simoa)^33^, an ultrasensitive bead-based immunoassay^34^ (Schematic 1B). Together, these techniques allow us to quantify and characterise tau aggregates with high sensitivity and molecular specificity, without the need for conformation-specific antibodies. This enables not only detection of aggregate levels, but also detailed profiling of their size, morphology and post-translational modifications. In parallel, we assessed the seeding activity of sarkosyl-insoluble tau aggregates using isoform-specific biosensor assays, to explore potential functional consequences (Schematic 1C).

Our goal was to determine whether the small diffusible tau aggregates show disease-specific features, and whether their molecular characteristics could shed light on the distinct pathological mechanisms of different tauopathies.

## RESULTS

### Quantitative profiling of tau species reveals disease-specific aggregation patterns

We began by broadly characterising tau species and their distribution across tauopathies using post-mortem brain homogenate from individuals with pathologically confirmed and clinically concordant AD (AD dementia), PSP (PSP Richardson’s syndrome or PSP-Parkinsonism), CBD (corticobasal syndrome), PiD (frontotemporal dementia), and age-matched neurologically healthy controls (HC) with mild age-related pathology. We measured the amounts of tau and tau aggregates in the brain homogenate fraction, and sarkosyl insoluble tau, as well as their relative distributions across both fractions. All samples were derived from the same brain region (frontal cortex) to enable direct comparison. Summarised patient details are listed in Table 1 (full details in Table 2).

**Table 1:**
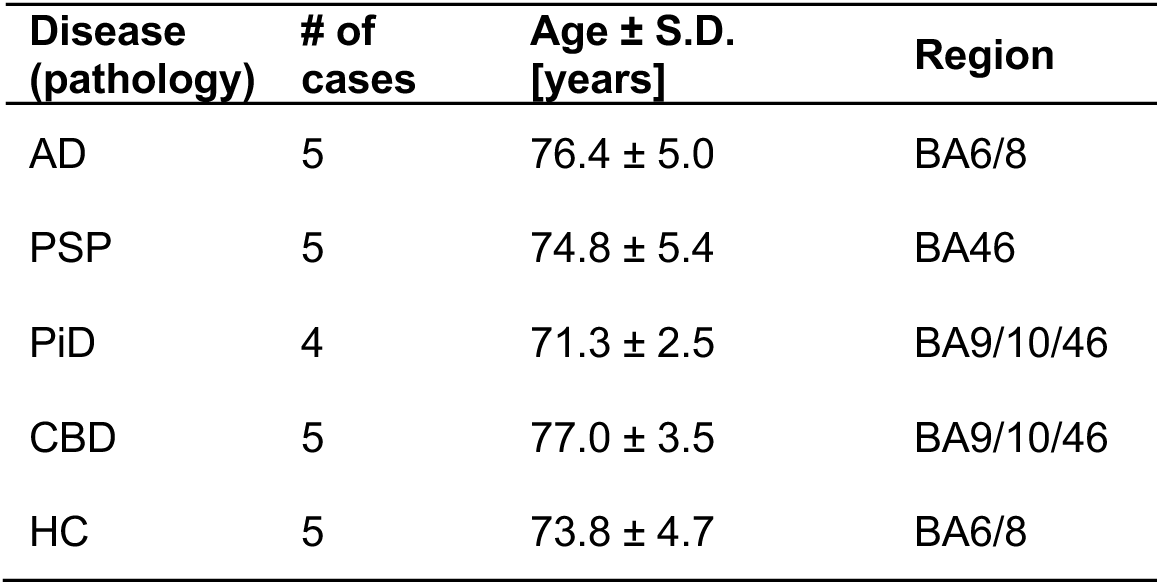
Summarised patient characteristics.

| **Disease (pathology)** | **# of cases** | **Age ± S.D. [years]** | **Region** |
| --- | --- | --- | --- |
| AD | 5 | 76.4 ± 5.0 | BA6/8 |
| PSP | 5 | 74.8 ± 5.4 | BA46 |
| PiD | 4 | 71.3 ± 2.5 | BA9/10/46 |
| CBD | 5 | 77.0 ± 3.5 | BA9/10/46 |
| HC | 5 | 73.8 ± 4.7 | BA6/8 |

**Table 2:**
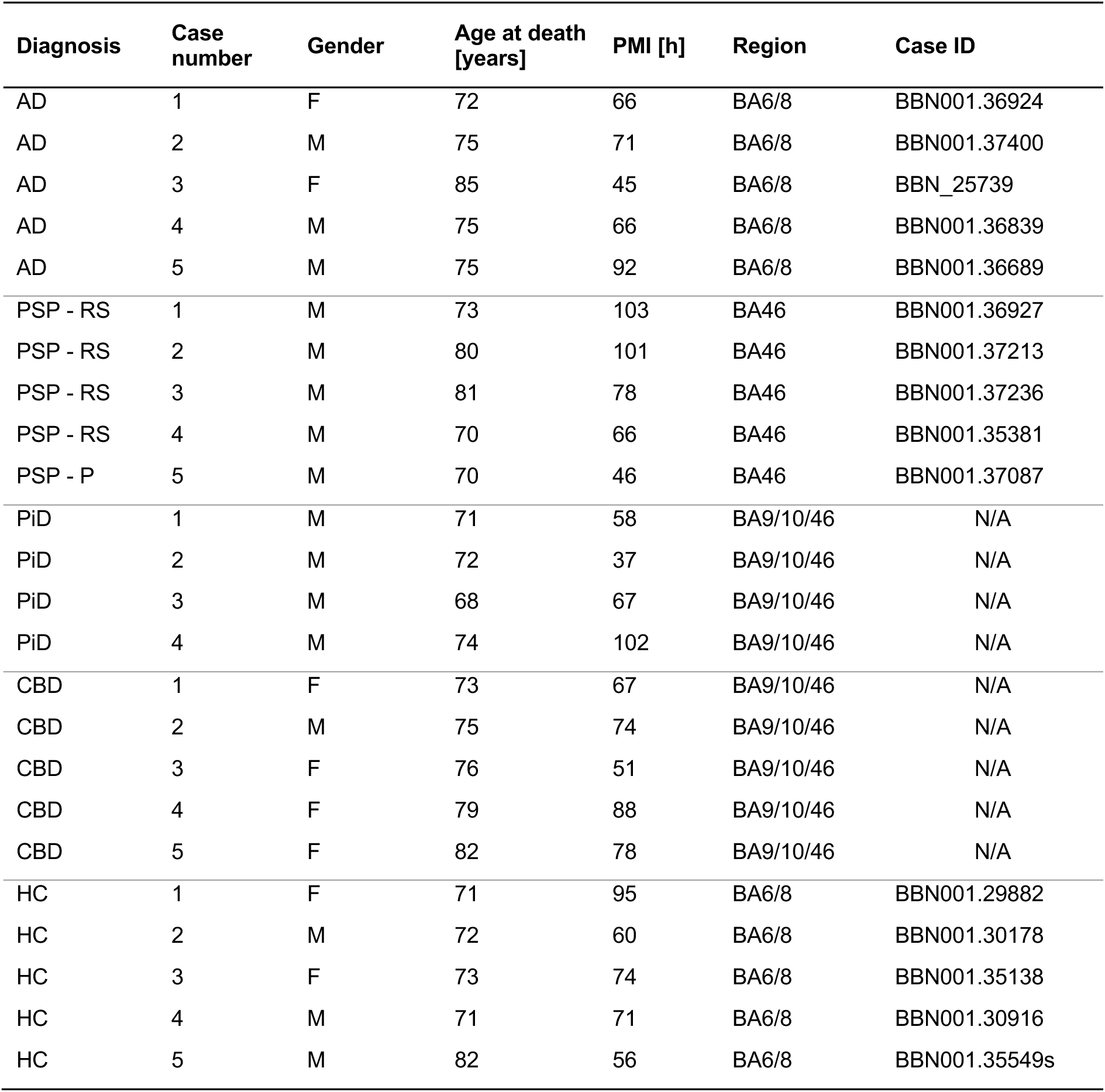
Detailed patient characteristics. RS: Richardson Syndrome, P: Parkinsonism-type presentation

| Diagnosis | Case number | Gender | Age at death [years] | PMI [h] | Region | Case ID |
| --- | --- | --- | --- | --- | --- | --- |
| AD | 1 | F | 72 | 66 | BA6/8 | BBN001.36924 |
| AD | 2 | M | 75 | 71 | BA6/8 | BBN001.37400 |
| AD | 3 | F | 85 | 45 | BA6/8 | BBN_25739 |
| AD | 4 | M | 75 | 66 | BA6/8 | BBN001.36839 |
| AD | 5 | M | 75 | 92 | BA6/8 | BBN001.36689 |
| PSP - RS | 1 | M | 73 | 103 | BA46 | BBN001.36927 |
| PSP - RS | 2 | M | 80 | 101 | BA46 | BBN001.37213 |
| PSP - RS | 3 | M | 81 | 78 | BA46 | BBN001.37236 |
| PSP - RS | 4 | M | 70 | 66 | BA46 | BBN001.35381 |
| PSP - P | 5 | M | 70 | 46 | BA46 | BBN001.37087 |
| PiD | 1 | M | 71 | 58 | BA9/10/46 | N/A |
| PiD | 2 | M | 72 | 37 | BA9/10/46 | N/A |
| PiD | 3 | M | 68 | 67 | BA9/10/46 | N/A |
| PiD | 4 | M | 74 | 102 | BA9/10/46 | N/A |
| CBD | 1 | F | 73 | 67 | BA9/10/46 | N/A |
| CBD | 2 | M | 75 | 74 | BA9/10/46 | N/A |
| CBD | 3 | F | 76 | 51 | BA9/10/46 | N/A |
| CBD | 4 | F | 79 | 88 | BA9/10/46 | N/A |
| CBD | 5 | F | 82 | 78 | BA9/10/46 | N/A |
| HC | 1 | F | 71 | 95 | BA6/8 | BBN001.29882 |
| HC | 2 | M | 72 | 60 | BA6/8 | BBN001.30178 |
| HC | 3 | F | 73 | 74 | BA6/8 | BBN001.35138 |
| HC | 4 | M | 71 | 71 | BA6/8 | BBN001.30916 |
| HC | 5 | M | 82 | 56 | BA6/8 | BBN001.35549s |

The small (soluble) tau fraction was obtained from a high-salt, low-detergent homogenisation buffer (10 mM Tris-HCl, 0.8 M NaCl, 1 mM EGTA, 0.1% Sarkosyl, 10% sucrose, 21,000 × g), which retains small, diffusible tau species^32^. Within this fraction (“homogenate”), we first measured total tau (monomeric and aggregated tau) using a commercial sandwich ELISA targeting pan tau. While CBD, PiD, AD, and HC showed similar levels of tau, PSP exhibited significantly lower levels (Figure 1A).

**Figure 1:**
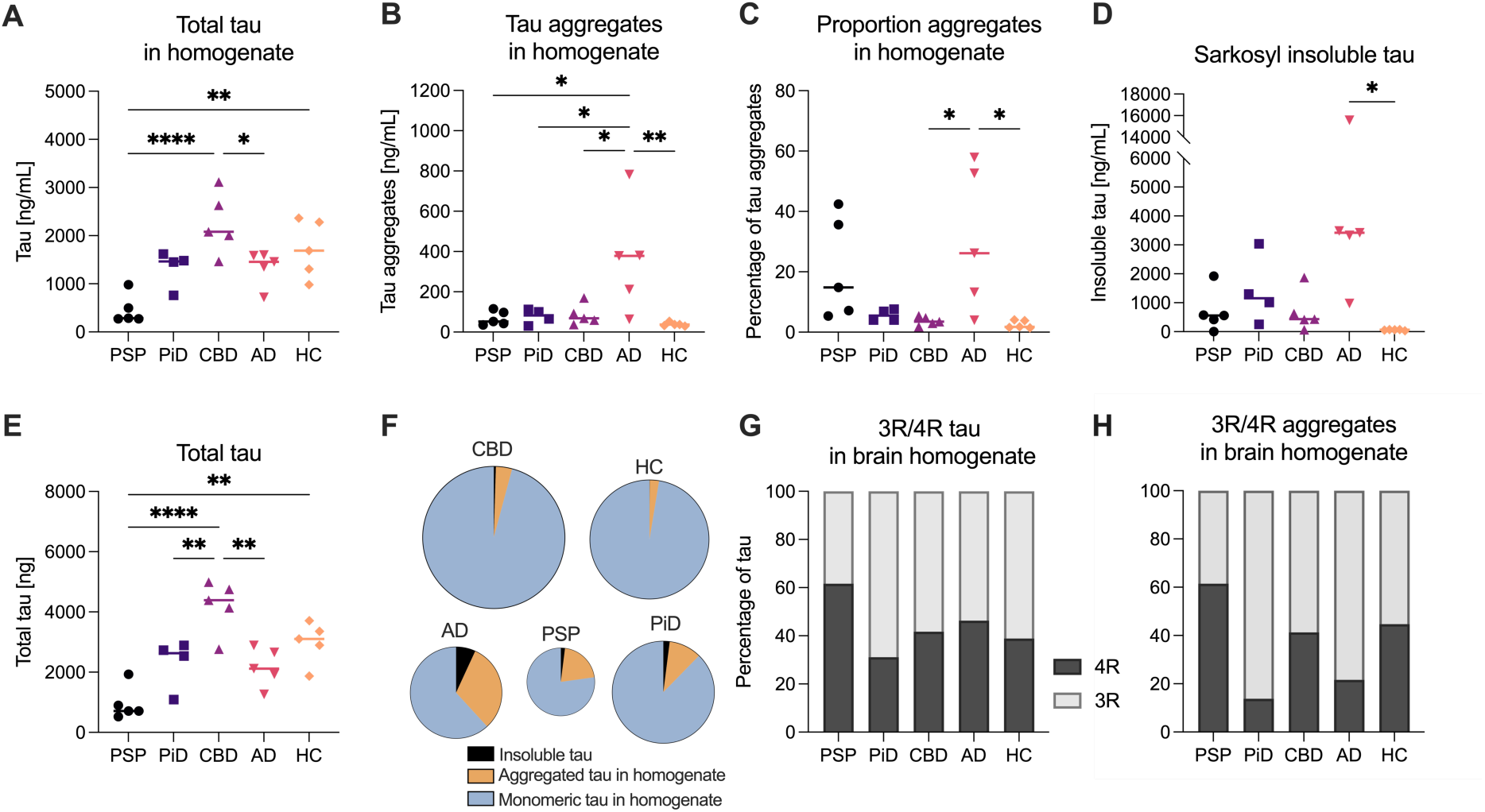
Differential distribution of tau species across tauopathies. (A) Total tau levels measured in brain homogenate from individuals with PSP, PiD, CBD, AD, and age-matched healthy controls (HC) using a mid-region tau ELISA. (B) Small tau aggregate levels measured in the same fraction using a single-molecule Simoa assay with an HT7 antibody sandwich configuration. (C) Ratio of small aggregates to total tau in brain homogenate. (D) Sarkosyl insoluble tau levels measured by ELISA in a separate sarkosyl extraction. (E) Total tau levels (brain homogenate + insoluble) across groups. (F) Relative distribution of tau species (monomer, small aggregates, and sarkosyl-insoluble tau) as a fraction of total tau for each disease group. The size of the pie charts reflects the total amount of tau. (G) Distribution of 3R and 4R tau isoforms in the total tau pool in brain homogenate, measured using a 4R-specific ELISA and normalised to total tau). (H) Isoform composition of small tau aggregates measured using a 4R-specific tau aggregate Simoa assay. In (A) - (E), each data point represents the mean of duplicate measurements of the individual cases (n = 4 cases for PiD, n = 5 for other disease group); horizontal lines indicate group medians. Statistical comparisons were performed using one-way ANOVA with Tukey’s post hoc test; * p < 0.05, **p < 0.01, ****p < 0.0001. For (F) – (H), the average of the n = 5 cases (n = 4 cases for PiD) was used. For clarity, only statistically significant comparisons are shown.

To quantify the small tau aggregates within the brain homogenate fraction, we applied a previously developed Simoa assay using an HT7 antibody sandwich configuration, which selectively detects aggregated tau^33^ and not monomers. Small tau aggregate levels were significantly elevated in AD brain homogenate compared to all other disease groups and HC (Figure 1B). When normalised to total tau levels in the homogenate, the proportion of tau present as aggregates was the highest in AD, but notably also elevated in PSP (Figure 1C). In contrast, CBD and HC showed similarly low aggregate-to-total tau ratios.

To further assess the distribution of tau into monomeric, small aggregates and insoluble species, we performed a separate sarkosyl extraction to isolate the detergent-insoluble tau fraction. We quantified insoluble tau aggregates (Figure 1D) and determined the absolute total amount of tau aggregates (soluble tau and sarkosyl insoluble tau, in ng, extracted from 120 mg of post-mortem brain) (Figure 1E). We also compared the relative contribution of monomeric and aggregated tau in the brain homogenate, and sarkosyl insoluble tau (Figure 1F, Supplementary Figure S1A). Due to epitope availability particularly in the insoluble fraction their amount is likely underestimated. AD showed the highest levels of sarkosyl-insoluble tau, with lower levels in CBD, PiD and PSP, while HC had minimal levels. When comparing the relative contribution of each tau species to the total tau pool, striking differences emerged. In AD, a substantial portion of total tau (7 ± 6%) was present as insoluble aggregates, whereas in CBD only a small fraction was insoluble (0.4 ± 0.4%), despite high overall tau levels (Figure 1E, F). This suggests that CBD maintains a large pool of tau with limited conversion into aggregated forms, while PSP, with low total tau levels, shows a relatively high proportion of aggregation in the brain homogenate fraction.

As the tauopathies included in this study are characterised by distinct tau isoform distributions (PiD is classified as a 3R tauopathy, PSP and CBD as 4R tauopathies, and AD as a mixed 3R/4R tauopathy) we next assessed isoform composition within the tau pool in the brain homogenate. We quantified 4R tau levels using a commercial ELISA, and developed an additional Simoa assay that specifically detects 4R-positive aggregates (Supplementary Figure S1B, C), which we normalised to total tau and total tau aggregate levels in the brain homogenate, respectively (Figure 1G, H, Supplementary Figure S1D). These ratios are approximate since some 4R aggregates also contain 3R tau. The resulting 3R/4R patterns in both total tau and aggregates in the brain homogenate were overall consistent with the known disease classifications, indicating that isoform-specific aggregation is preserved in the soluble phase.

### Super-resolution imaging reveals disease-specific tau aggregate morphologies

Having established that the small tau aggregates present in brain homogenate differ in abundance and distribution across tauopathies, we next characterised their morphology, a property that remains largely unexplored but may provide insight into disease-specific aggregate features. We applied single-molecule super-resolution microscopy to tau aggregates captured from brain homogenate using the MAPTau^32^. Aggregates were immobilised on a PEG coated surface using monoclonal tau antibodies, labelled with the same monoclonal, fluorescently conjugated antibody, and imaged with TIRF microscopy, enabling visualisation and quantitative analysis of individual aggregates. For each aggregate, we extracted morphological parameters including length, area, and eccentricity to compare the aggregate shape and size distributions across diseases.

Representative super-resolution images revealed small, round aggregates in all groups, while AD samples also contained a distinct population of long (>200 nm), fibrillar-shaped aggregates not observed in other tauopathies or HC (Figure 2A). To quantify this, we first measured the mean aggregate length per sample, which was significantly higher in AD compared to PSP, PiD, CBD, and HC (Figure 2B). The cumulative distributions of aggregate length showed a right-shifted profile in AD consistent with a higher proportion of longer aggregates (Figure 2C). Importantly, the presence of long aggregates in AD was not attributable to overall aggregate abundance: they remained detectable even upon dilution, while PSP samples imaged at fivefold higher concentration remained crowded but exclusively composed of small, round aggregates (Supplementary Figure S1E).

**Figure 2:**
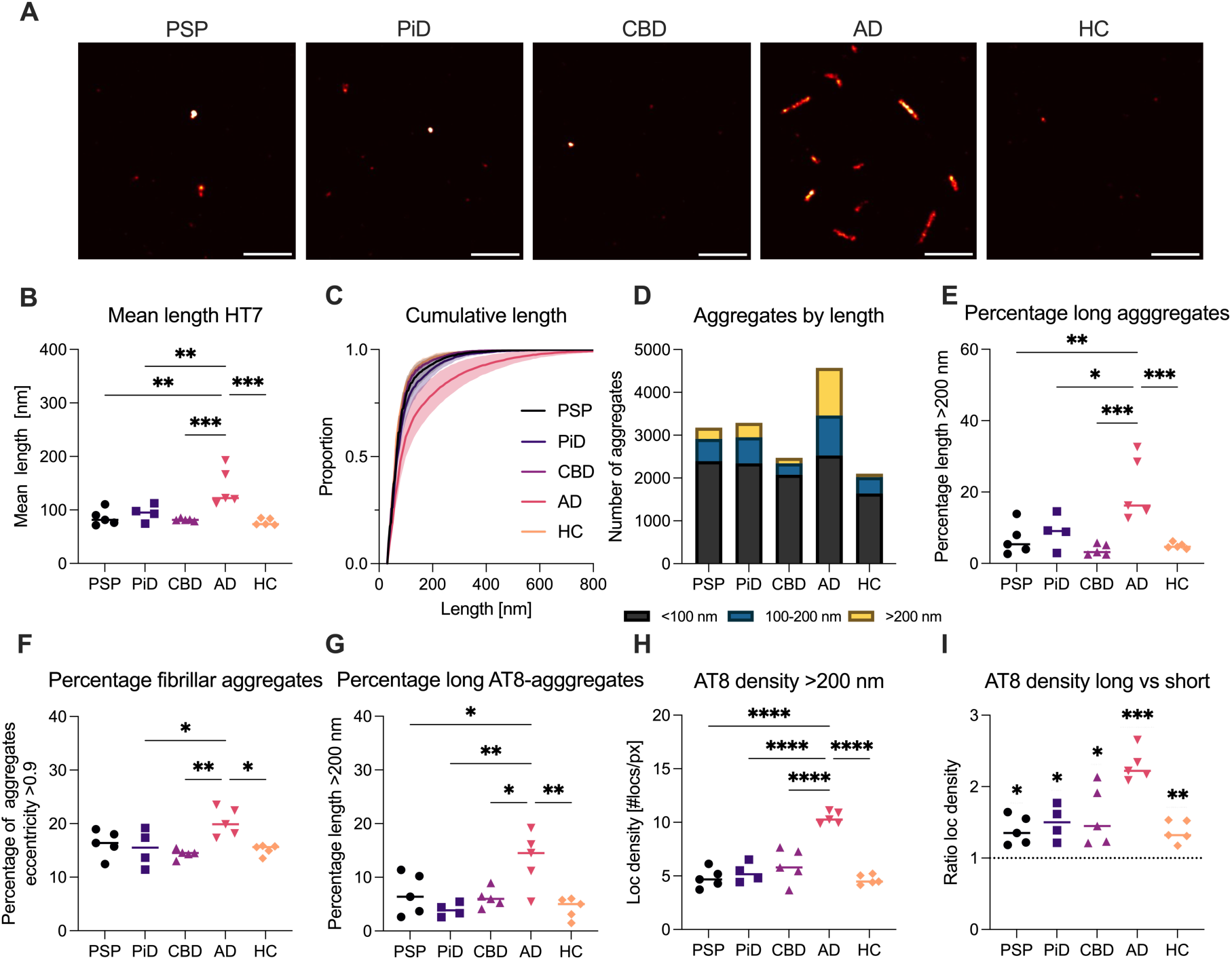
Super-resolution imaging reveals disease-specific tau aggregate morphologies. (A) Representative single-molecule super-resolution images of small tau aggregates in the brain homogenate fraction captured with HT7 antibodies from brain homogenate. Scale bar, 1 µm. (B) Quantification of mean aggregate length. (C) Cumulative distribution of aggregate length. Shown is mean individual cases for each cohort, shading depicts ± S.D.. (D) Total number of aggregates per group, categorised by length into <100 nm (black), 100–200 nm (blue), and >200 nm (yellow). (E) Proportion of aggregates with length >200 nm, used to define “long” aggregates. (F) Proportion of aggregates with eccentricity >0.9, indicative of a fibrillar morphology. (G) Proportion of AT8-positive aggregates with length >200 nm. (H) Localisation density (localisations per pixel) of AT8-positive aggregates >200 nm in length. (I) Ratio of localisation density in long (>200 nm) versus short (<100 nm) AT8-positive aggregates for each sample. In (B) - (D) and (F) – (I), each data point represents an individual case (n = 4 for PiD, n = 5 per other disease group); horizontal lines indicate group medians. Statistical comparisons were performed using one-way ANOVA with Tukey’s post hoc test, or one-sample t-test in panel (I); \**p* < 0.05, **p < 0.01, ***p < 0.001, ****p < 0.0001. For (E), the mean of the n = 5 cases was used (n = 4 for PiD).

To better classify aggregate length distributions across diseases, we categorised all aggregates by length into three bins: small (<100 nm), medium (100–200 nm), and long (>200 nm) (Figure 2D). The number of small aggregates was comparable across all tauopathies (2335 ± 189 aggregates per FOV) and slightly lower in healthy controls, indicating a shared baseline of early aggregation. In contrast, AD exhibited an increase in both medium and long aggregates, suggesting the presence of an additional, disease-specific population. When expressed as a proportion, long aggregates (>200 nm) were significantly more abundant in AD (21 ± 8.9%) compared to other tauopathies (6.5 ± 3.9%) (Figure 2E). Consistently, AD also showed a higher proportion of elongated, fibrillar-shaped aggregates (eccentricity >0.9; AD: 21 ± 2%, other tauopathies: 15 ± 2%; Figure 2F).

To assess whether these fibrillar-shaped aggregates also carried disease-relevant post-translational modifications, we performed super-resolution imaging using the phospho-tau specific AT8 antibody (pS202/pT205). As with HT7, AT8-positive aggregates in AD also included long fibrillar-shaped species, and the proportion of AT8-positive aggregates >200 nm was significantly higher in AD than in every other group (Figure 2G, mean length and percentage of fibrils of AT8-positive tau aggregates in Supplementary Figure S1F).

We next assessed AT8 epitope availability by calculating the localisation density (localisations per super-resolved pixel, corresponding to 13x13 nm^2^) within each aggregate as a proxy for phosphorylation load. In long aggregates (>200 nm), AD showed significantly increased AT8 localisation density compared to all other groups (Figure 2H), with a mean of 10.4 ± 0.5 localisations/pixel versus 5.5 ± 1.4 localisations/pixel in the other conditions. This difference was specific to AT8, as HT7-based localisation densities were similar across aggregate lengths and disease states (Supplementary Figure S1G). Finally, to compare phosphorylation between short and long aggregates, we calculated the localisation density ratio (>200 nm vs. <100 nm). All groups showed enrichment of AT8 in long aggregates (Figure 2I), suggesting this modification is generally associated with long, fibrillar forms.

Together, these findings indicate that AD is characterised by a specific population of long, fibrillar-shaped tau aggregates in the brain homogenate fraction, which may be structurally and biochemically distinct from those in other tauopathies.

### Small tau aggregates exhibit disease-specific phosphorylation profiles

Given the morphological and AT8 phosphorylation status differences observed in AD, we next sought to systematically examine the phosphorylation patterns of small tau aggregates across tauopathies. To do this, we employed aggregate-specific Simoa assays targeting defined phospho-epitopes, alongside single-molecule co-localisation microscopy, to map the presence and overlap of key phosphorylation sites on individual tau aggregates.

We first quantified the overall abundance of small aggregates in the brain homogenate that carry specific phospho-epitopes using aggregate-specific Simoa assays. We used the previously developed assay to detect AT8 (pS202/pT205) positive tau aggregates and developed further assays to specifically detect tau aggregates positive for pT217, pT181, and pS356 rather than monomers (Supplementary Figure S2A). AD samples showed significantly increased levels of AT8-, pT217-, and pT181-positive aggregates compared to the other tauopathies and controls (Figure 3A–C). To account for differences in overall aggregate burden, we normalised to the levels of total tau aggregates in the brain homogenate fraction as detected by the HT7 antibody pair. Phosphorylation at AT8 and T217 remained significantly enriched in AD, while T181 no longer showed significant differences across groups (Supplementary Figure S2B-E). In contrast, aggregates positive for phosphorylation at S356 showed a distinct profile. pS356-positive aggregates were most abundant in PSP and nearly absent in AD (Figure 3D). Total pS356 levels did not differ across diseases (Supplementary Figure S2F), indicating that pS356 is present in AD but likely only in monomeric tau or masked within aggregates, suggested by recent findings^35^.

**Figure 3:**
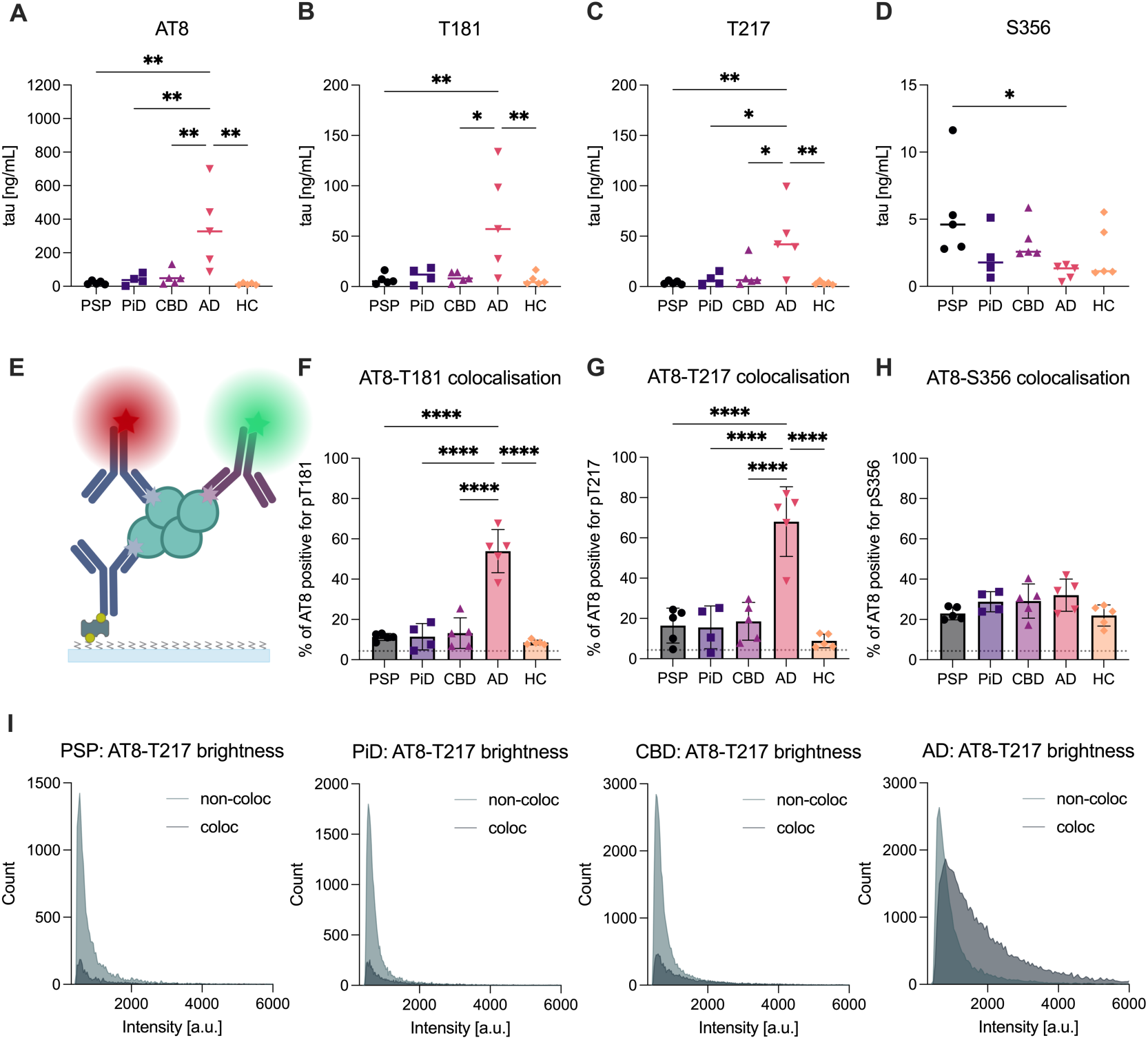
**Small tau aggregates exhibit disease-specific phosphorylation profiles**. (A–D) Aggregate-specific Simoa assays quantifying small tau aggregates positive for (A) AT8 (pS202/pT205), (B) pT217, (C) pT181, and (C) pS356. (E) Schematic of two-colour single-molecule co-localisation microscopy used to determine co-occurrence of phospho-epitopes. (F–H) Proportion of AT8-positive aggregates co-labelled with (F) pT217, (G) pT181, or (H) pS356, where the dotted-line indicates the by-chance-co-localisation. (I) Fluorescence intensity distributions of AT8-positive aggregates stratified by co-labelling status for each phospho-epitope. Each point represents one individual with n = 4 cases for PiD and n = 5 cases for other disease groups. Bars show mean ± SD. Statistical comparisons were performed using one-way ANOVA with Tukey’s post hoc test; \**p* < 0.05, **p < 0.01, ***p < 0.001, ****p < 0.0001.

To assess the co-occurrence of phospho-epitopes within individual aggregates, we performed two-colour single-molecule co-localisation imaging (Figure 3E, Supplementary Figure S2G-I for 488) and quantified the proportion of AT8-positive aggregates that also carried a second phospho-epitope (T217, T181, or S356). In AD, on average 54 ± 11% of AT8-positive aggregates were co-labelled with pT181 and 68 ± 17% with pT217. In contrast, less than 20% of AT8-positive aggregates were positive for either epitope in PSP, CBD, and PiD, and fewer than 10% showed co-localisation in HC samples (Figure 3F, G). For S356, co-localisation with AT8 was uniform across all groups (around 30%, Figure 3H), consistent with its distinct aggregate profile.

We next analysed the fluorescence intensity of individual aggregates, using signal intensity as a proxy for aggregate size and/or epitope density. In AD, AT8-positive aggregates were overall brighter, consistent with the increased size/phosphorylation density observed by super-resolution imaging (pT217 Figure 3I, pT181 and pS356 in Supplementary Figure S2J). Aggregates were then stratified based on whether they were co-labelled with a second phospho-epitope (pT181, pT217 or pS356). Co-localised aggregates were overall brighter/larger, with most high-intensity aggregates showing co-localisation. In PiD and CBD, AT8 aggregates were generally dimmer, consistent with their smaller size as observed by super-resolution imaging. Both co-labelled and non-co-labelled aggregates were present among the brighter population. In contrast, PSP lacked bright co-labelled aggregates entirely, with high-intensity aggregates observed only in the non-co-labelled fraction.

Together, these results show that the small tau aggregates differ not only in their phosphorylation levels but also in the co-occurrence of specific phospho-epitopes at the single-aggregate level.

### Ubiquitination and C-terminal truncation distinguish aggregate subtypes across tauopathies

To further investigate the disease-specific differences in aggregate composition, we examined additional post-translational modifications, specifically ubiquitination, and probed the availability of N-and C-terminal tau epitopes to assess potential truncation.

AD showed the highest proportion of AT8-positive aggregates co-labelled with ubiquitin, while other tauopathies and controls displayed markedly lower levels of ubiquitin co-labelling (Figure 4A and Supplementary Figure 3A, B). Across all groups, ubiquitin-positive aggregates were generally brighter than non-ubiquitinated ones (measured by AT8 brightness) (Supplementary Figure S2J), suggesting that ubiquitinated aggregates tend to be larger.

**Figure 4:**
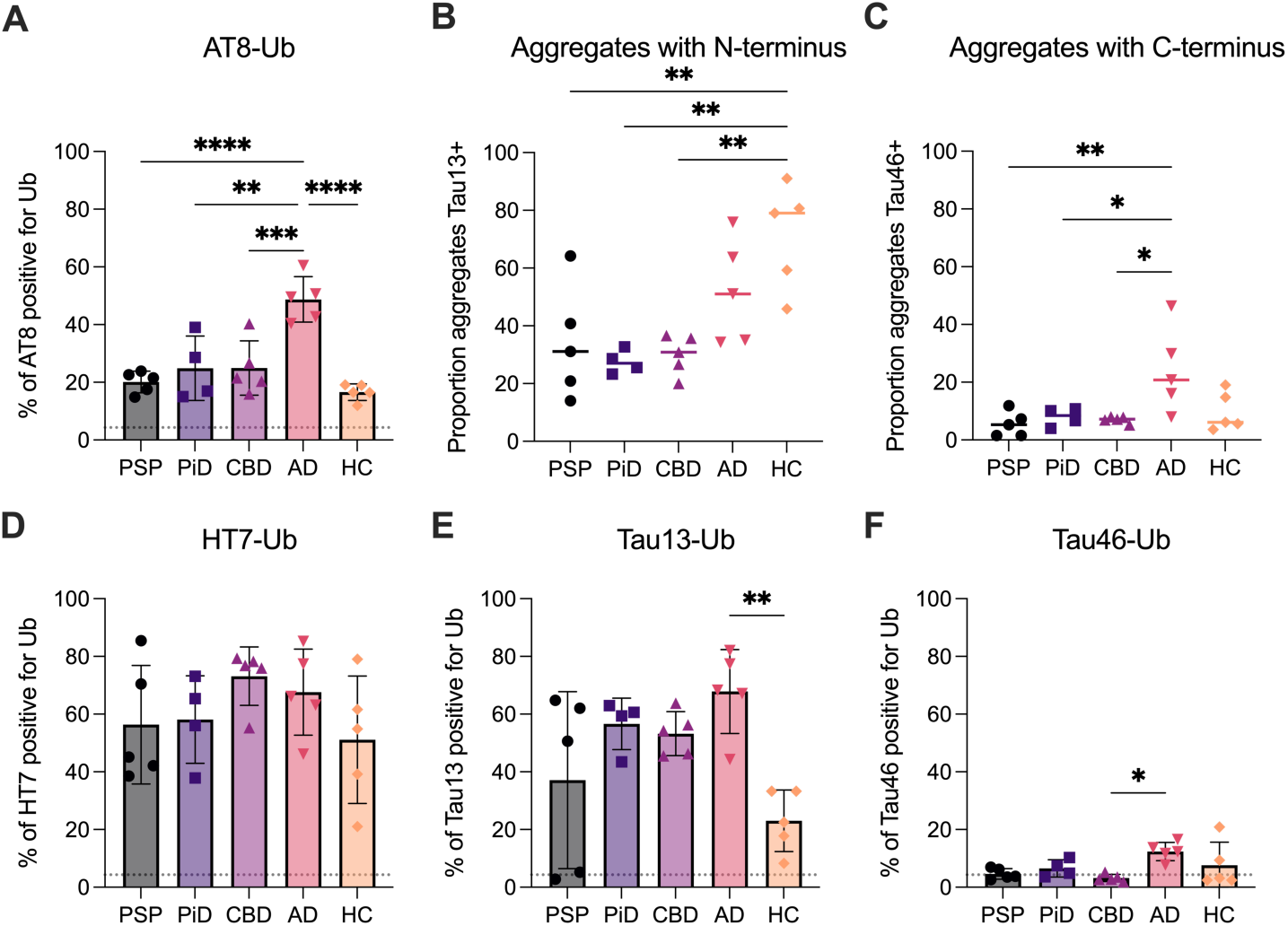
**Disease-specific patterns of tau ubiquitination and truncation**. (A) Proportion of AT8-positive tau aggregates co-labelled with ubiquitin determined by two-colour single-molecule imaging. (B–C) Proportion of tau aggregates retaining the (B) N-terminal epitope (Tau13) and (C) C-terminal epitope (Tau46), normalised to total tau aggregate levels (HT7), determined through Simoa. (D–F) Proportion of (D) HT7-, (E) Tau13-, and (F) Tau46-positive tau aggregates with ubiquitin, determined by two-colour single-molecule imaging. Dotted line represents the by chance co-localisation. Each data point represents one individual (n = 4 for PiD, n = 5 for other disease groups); bars mean ± SD. Statistical comparisons were performed using one-way ANOVA with Tukey’s post hoc test; \**p* < 0.05, \*\**p* < 0.01, \*\*\**p* < 0.001, **** *p* < 0.0001.

To assess whether tau truncation also differed between diseases, we probed the availability of N-and C-terminal tau epitopes using aggregate-specific Simoa assays (Tau13 detecting N-terminus, and Tau46 detecting the C-terminus; thereby detecting aggregates containing at least two monomers with the respective terminus). We normalised these to the levels of total tau aggregates determined through HT7 (mid-region epitope), to determine the proportion of truncated aggregates. A large proportion of tau aggregates retained the N-terminus across all diseases (Figure 4B). However, the C-terminus was detected in less than 10% of aggregates in controls and non-AD tauopathies, while in AD, on average 22 ± 12% aggregates retained the C-terminus (Figure 4C). This suggests that C-terminal truncation is widespread in the small tau aggregates in non-AD tauopathies but less frequent in AD.

We next examined co-localisation of each tau epitope (HT7, Tau13, Tau46) with ubiquitin using two-colour single-molecule imaging. HT7 and Tau13 aggregates showed moderate to high levels of ubiquitin co-labelling across all disease groups (Figure 4D, E, HT7-Ub: 62 ± 17%, Tau13-Ub: 45 ± 24%). In contrast, Tau46-positive aggregates rarely co-localised with ubiquitin (Figure 4F, Tau46-Ub: 6.7 ± 5.0%). This indicates that aggregates containing at least two tau proteins with intact C-terminus are less likely to be ubiquitinated, suggesting a potential mutual exclusivity between C-terminal truncation and ubiquitin tagging.

### Small tau aggregates exhibit disease-distinct molecular signatures

To visualise these multidimensional differences, we generated a radar plot showing the average profile of each disease across key features (Figure 5A). This highlights the distinct molecular “fingerprints” of the small tau aggregates in each condition, with AD showing a particularly divergent profile driven by aggregate length, phosphorylation, and ubiquitination in the frontal cortex.

**Figure 5:**
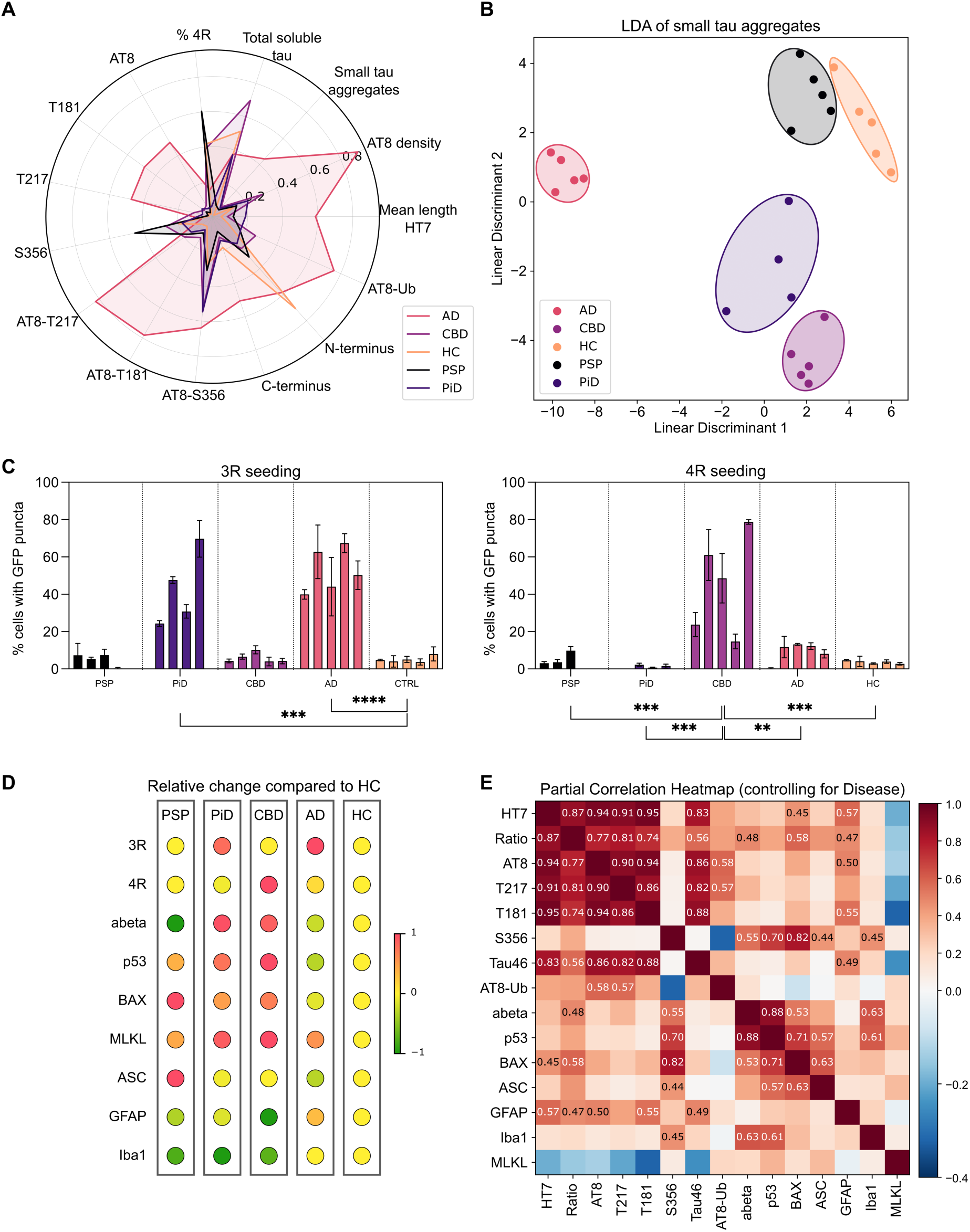
Small tau aggregates encode disease-specific molecular profiles and divergent functional signatures. (A) Radial distribution plot of small tau aggregate features (scaled within each feature. (B) Linear discriminant analysis (LDA) of small tau aggregate features. LD1 explains 79.6% of between-class variance while LD2 explains 14.0% of variance. (C) Isoform-specific tau seeding activity in HEK293 biosensor cell lines expressing either 3R-or 4R-tau. Cells were seeded with equal amounts of sonicated sarkosyl-insoluble tau and the percentage of seeded cells was quantified after 72h based on eGFP puncta. Bars show the mean ± S.D. of the n = 3 technical replicates for each individual (n = 4 for PiD, n = 5 per other disease group). (D) Relative levels of selected stress and inflammatory markers across tauopathies, normalised to the mean of HC samples. Markers include p53 (aggregated form, measured via Simoa), BAX and MLKL (apoptosis and necroptosis, measured via ELISA), GFAP and Iba1 (astrocytic and microglial activation, measured via ELISA), and ASC specks (inflammasome activation, measured via SiMPull). Values are scaled from –1 to 1 to allow cross-marker comparison. (E) Partial correlation matrix showing associations between tau aggregate features and stress or inflammatory markers, controlling for disease group. Only significant associations (p<0.05) are shown. Values indicate partial correlation coefficients, controlling for disease group.

Having characterised individual features of the small tau aggregates across tauopathies, we next asked whether these small tau aggregates differ systematically between diseases, i.e., if the features of these aggregates can be utilised in combination to categorise different tauopathies. While disease-specific folds of insoluble tau are well-established, it remains unclear whether this is the case for the earlier-stage, small tau aggregates.

To address this, we performed linear discriminant analysis (LDA) using only measured small tau aggregate features, including abundance, morphology, phosphorylation, and ubiquitination. This revealed clear separation of disease groups in multivariate space, indicating that the composition and structural properties of the small tau aggregates differ across tauopathies (Figure 5B). Linear discriminant 1 (LD1), with percentage of long aggregates, C-and N-terminal truncation as top contributing features, explained 69.0% of between-class variance while LD2 explained 22.3% of variance, with percentage of long aggregates, percentage of 4R tau aggregates and pS356 as top contributing features. Notably, AD samples clustered furthest from all other groups on the first dimension, and CBD on the second dimension, consistent with their distinct fibrillar aggregate morphology and PTM profile. Together, this suggests that small tau aggregates encode disease-specific molecular information.

### Functional and cellular signatures associated with tau aggregates

#### Divergent seeding profiles and cell stress marker expression across diseases

Building on our findings that the small tau aggregates differ in their characteristics across tauopathies, we next explored whether these differences were accompanied by distinct functional properties and associated tissue-level stress markers. While causality cannot be established from these measurements, such comparisons can provide insight into potential mechanisms of disease progression and divergent cellular responses.

### Isoform-specific seeding activity of sarkosyl-insoluble tau aggregates

While our preceding analyses focused on the distinct biochemical and structural properties of the small tau aggregates in the brain homogenate fraction, we next asked whether insoluble tau aggregates differ in their capacity to propagate tau pathology. Although the small aggregates characterised above are hypothesised to be toxic and potentially seed-competent, we observed no seeding above background when using homogenate directly, likely due to the high protein load impairing cell viability. As a result, we turned to sarkosyl-insoluble tau aggregates, which are enriched in misfolded conformers and are more commonly used in established biosensor assays. Seeding assays provide a functional, quantitative readout of tau’s ability to induce aggregation in a prion-like manner, a process suggested to underlie tau propagation and disease progression *in vivo*.

To evaluate isoform-specific seeding activity, we used tau biosensor HEK293 cell lines expressing either full-length wild-type 3R- or 4R-tau fused to eGFP^36^. The cells were seeded with approximately the same amount of sonicated sarkosyl insoluble tau (determined through ELISA), and the seeding efficiency was evaluated as the percentage of seeded cells based on eGFP puncta.

We observed distinct seeding patterns across diseases (Figure 5C). AD and PiD samples induced strong aggregation in the 3R biosensor line, consistent with their 3R or mixed 3R/4R tau pathology. CBD exhibited robust seeding in the 4R line, while PSP showed little to no seeding in either cell line, despite its 4R profile. AD also showed moderate seeding in the 4R line. Only background levels of seeding were observed in HC samples.

These results demonstrate that the seeding competence of insoluble tau aggregates varies across tauopathies and aligns with their isoform composition, suggesting disease-specific conformations and templating capacity.

To confirm the isoform composition of the sarkosyl-insoluble aggregates used in the seeding assays, we performed western blot analysis on the same samples using pan-tau, 3R-specific, and 4R-specific antibodies (Supplementary Figure 4A). The patterns were consistent with known isoform distributions in each disease: PiD showed predominant 3R-tau, PSP and CBD exhibited 4R-tau bands, and AD showed a mixture of both isoforms. These data validate the isoform-specific seeding responses observed in the biosensor assay and support the conclusion that tau aggregate conformations are disease-specific and isoform-dependent.

### Amyloid-β burden and markers of cell stress and gliosis

To explore additional pathological features that may accompany disease-specific tau profiles, we next quantified amyloid-β (Aβ) levels and a panel of markers related to cell stress, death pathways, and gliosis (relative to HC in Figure 5D, without normalisation in Supplementary Figure S4B-H).

Aβ aggregate burden was assessed using a previously established Simoa aggregate assay employing the 6E10 antibody^33^. No significant differences were observed across disease groups, suggesting a largely age-related background signal in these samples.

We measured several markers of cell death and inflammatory responses (Figure 5D), including p53 aggregates, a stress-responsive transcription factor involved in DNA repair, as well as programmed cell death^37,38^ (measured through Simoa^39^). Using commercially available ELISAs, we assessed BAX, a pro-apoptotic effector downstream of p53; MLKL, a key effector of necroptosis; and GFAP and Iba1 as markers of astrogliosis and microglial activation, respectively. Lastly, to capture inflammasome activation we measured ASC speck levels through SiMPull^40^.

Distinct patterns emerged across diseases. CBD exhibited high levels of p53, BAX, and MLKL, suggesting activation of both apoptotic and necroptotic responses. PiD similarly showed elevated BAX and p53, along with increased MLKL. PSP, by contrast, displayed high BAX but low MLKL, indicating a predominantly apoptotic profile. AD showed comparatively low BAX and p53 levels but increased Iba1 and GFAP, consistent with a more glial-dominant phenotype. ASC specks were most abundant in PSP and lower in all other groups, including AD.

These findings suggest that tauopathies not only differ in tau aggregation and phosphorylation profiles, but also in the pattern of associated inflammatory and cell stress responses, with variable involvement of apoptotic, necroptotic, and glial pathways.

### Correlation analysis links aggregate markers to inflammatory and cell stress responses

To summarise these disease-specific patterns and facilitate comparison across tauopathies, we generated a matrix showing relative marker changes compared to HC samples (Figure 5D). This matrix highlights distinct cellular stress and inflammatory profiles for each disease, reinforcing the concept that despite overlapping tau pathology, each tauopathy features unique pathological and cellular signatures.

To explore potential relationships between tau aggregate characteristics and cellular stress markers, we performed a partial correlation analysis across all measured parameters, controlling for disease group (Figure 5E, correlation coefficient only shown for p < 0.05). These included aggregate-related features such as phosphorylation sites, truncation, and ubiquitination, alongside markers of inflammation, cell death, and Aβ aggregate burden.

As expected, p53 and BAX were strongly correlated, consistent with their known regulatory relationship in apoptotic signalling^37^. p53 also showed moderate correlations with ASC specks, Iba1, and Aβ, suggesting a broader stress response involving both proteinopathy and inflammation.

Among tau-related features, pS356-positive tau aggregates, despite lacking correlation with other tau phosphorylation sites, displayed the most extensive associations with other markers. For example, pS356-positive aggregates showed a strong correlation with BAX (r = 0.81), and moderate correlations with p53, ASC, Iba1, and Aβ (p53: r = 0.65, ASC: r = 0.46, Iba1: r = 0.46, Aβ: r = 0.53). These findings suggest that phosphorylation at S356 may mark a subset of tau aggregates associated with apoptotic and inflammatory signalling, particularly relevant in diseases such as PSP where this site is enriched.

Additional associations included a moderate correlation between GFAP and the total tau aggregate levels as well as AT8 and T181-positive aggregates, potentially linking hyperphosphorylated tau species to astrogliosis, especially in AD. The proportion of tau aggregates showed moderate correlation with BAX, ASC and GFAP, indicating that high tau aggregation propensity and/or impaired clearance mechanism may be associated with cellular stress and inflammation.

Together, these data suggest that specific features of the small tau aggregates in brain homogenate, particularly PTMs, may co-vary with inflammatory and cell death responses in a disease-specific manner. While causality cannot be inferred, these correlations may reflect parallel pathological processes or shared upstream stressors shaping the molecular landscape of each tauopathy.

## DISCUSSION

The tauopathies are clinically and biologically heterogeneous neurodegenerative diseases, unified by the pathological aggregation of the tau protein. While much of the field has focused on large, insoluble fibrillar aggregates, increasing evidence suggests that small tau aggregates formed during the earlier stages of aggregation play a more central role in disease pathogenesis^21^. Here, we systematically characterised these small tau aggregates in brain homogenate across AD, PSP, CBD and PiD, using two complementary single-molecule techniques. We found that these aggregates differ not only in abundance but also in morphology, tau isoform composition, post-translational modifications, and associated markers of cellular stress. Together, our findings suggest that these small tau aggregates differ between diseases, with implications for molecular diagnostics and stratified therapies for tauopathies.

### Distinct aggregate properties reflect divergent disease mechanisms

A central finding of this study is that AD contains a distinct population of long, fibrillar-shaped aggregates in the brain homogenate fraction that are absent from other tauopathies. These fibrils are highly phosphorylated at disease-relevant sites (e.g. AT8, T217), and are associated with increased localisation density, suggesting a higher phosphorylation load per aggregate. In contrast, other diseases such as PSP and CBD displayed predominantly small, round aggregates, despite similar or higher levels of total tau in the brain homogenate fraction. This morphological divergence may reflect differences in tau conformation, proteostasis, or aggregation kinetics across diseases. That these fibrillar-shaped aggregates in AD are still diffusible yet structurally distinct supports the hypothesis that small tau aggregates may encode pathogenic characteristics before they transition to insoluble deposits.

Our findings further reveal that tau aggregates are not uniform across tauopathies, but rather vary in morphology, functional activity and associated cellular responses. Small aggregates exhibit disease-specific morphologies and post-translational modifications, which in turn correlate with differing biological effects in each disease. For example, in AD, fibrillar and hyperphosphorylated aggregates are accompanied by astroglial and microglial activation, while in CBD, tau aggregates show robust seeding activity but low associated inflammation. PSP presents a distinct profile: high proportion of small aggregates in the homogenate, low seeding, and high inflammasome activation, suggesting a localised, inflammatory pathology^42^. These observations suggest that tau aggregates may play different roles across diseases – acting as seeds, inflammatory triggers, or relatively inert by-products depending on their specific molecular features.

These interpretations are supported by recent findings showing that the morphology and PTMs of small tau aggregates differ across tauopathies and correlate with distinct cellular responses. Recent work supports the idea that not all small tau species are functionally equivalent. Meng *et al.* showed that hyperphosphorylated tau can spontaneously form small, round oligomers that trigger TLR4-dependent responses^28^. This provides a plausible mechanism by which the small, round aggregates could engage innate immune signalling. Yang *et al.* demonstrated that recombinant tau oligomers impair synaptic plasticity in a TNFα-dependent manner, whereas fibril-derived soluble aggregates disrupt plasticity independently of TNFα^41^.

In addition to morphology and phosphorylation, we also observed disease-specific patterns of tau ubiquitination and truncation. AD aggregates more frequently retained the C-terminal tau epitope and were less often ubiquitinated, whereas in non-AD tauopathies, aggregates lacking the C-terminus showed higher co-localisation with ubiquitin. This may suggest divergent aggregate clearance mechanisms or differential engagement of the ubiquitin-proteasome system across tauopathies, potentially contributing to differences in aggregate stability or turnover.

These findings reinforce the concept that distinct soluble tau aggregates have distinct functional properties, with some requiring or synergising with inflammatory cues to impair neuronal function. In this context, the combination of small tau aggregates and high inflammation observed in PSP may drive localised, rapid pathology^42^. In contrast, AD is less associated with inflammation in the frontal cortex and features more fibrillar-shaped aggregates, which may facilitate progressive spread, while CBD may follow a seeding-dominant but comparatively less inflammatory trajectory.

### pS356-positive aggregates as a distinct pathological subclass

Among all aggregate markers tested, phosphorylation at S356 stood out for its disease specificity and association with cellular stress. pS356-positive aggregates were enriched in PSP, nearly undetectable in AD, and strongly correlated with markers of apoptosis (BAX), inflammation (Iba1, ASC), and stress response (p53). Although total pS356 levels were similar across diseases, their reduced detectability in AD aggregates may reflect epitope burial, consistent with recent structural studies.

The structural accessibility of S356 is disease-dependent; while in PSP and CBD S356 is exposed on the outer surface of tau filaments, in AD PHFs it is buried within the fibril core. This suggests that pS356 not only marks 4R tauopathies but also reflects the conformational pathway underlying aggregation. Its presence on soluble aggregates in PSP and CBD indicates that even small tau species in these diseases adopt structures that permit phosphorylation at this site. In contrast, in AD, the inaccessibility of Ser356 may prevent such modification^35^ (even in the presence of relevant kinases), explaining the near absence of detectable pS356-positive soluble aggregates in AD in our study.

Given the known role of pS356 in blocking proteasomal degradation^43^, its presence in PSP may allow tau aggregates to evade clearance and accumulate, leading to the high proportion of soluble tau aggregates that we observed. While this could reflect an aberrant process or even a compensatory attempt to stabilise misfolded tau under stress, we suggest it more likely to ultimately lead to the build-up of bioactive tau species that may trigger inflammatory cascades. In support of this, we observed increased inflammatory and apoptotic markers in PSP, and specifically linked pS356-positive aggregates to elevated levels of BAX, p53, ASC specks, and Iba1. These inflammatory cascades may in turn further exacerbate tau pathology^44^. Additionally, the negative charge introduced at S356, located within the microtubule-binding domain, may also modulate local tau conformation and protein–protein interactions^45^, potentially altering the aggregation trajectory itself. Overall, these findings suggest that detectable S356 phosphorylation is both a structural and functional marker of tauopathies that adopt a PSP/CBD-like trajectory, with potential implications for disease classification and therapeutic targeting.

Our findings also raise the question of causality between PTMs and tau conformation. Do disease-specific PTMs, such as pS356, shape the aggregation trajectory and promote the adoption of particular filament folds? Or do distinct folds emerge early and constrain the accessibility of certain PTM sites, thereby encoding divergent modification profiles? The burial of sites such as S356 within the core of mature filaments, as seen in AD PHFs, suggests that once aggregation has occurred, conformational constraints may limit further modification, effectively locking in the PTM profile. However, in the early aggregation stages modifications may still play a key role in directing tau toward particular aggregation pathways. Understanding the temporal order of PTM acquisition and aggregation could provide crucial insight into the initiation and divergence of tauopathy subtypes.

### Fast vs slow tauopathy

Molecular and cellular consequences of disease-specific PTM profiles may also help explain the divergent patterns of tau aggregation observed across tauopathies. The higher abundance and fibrillar morphology of soluble tau aggregates in AD may reflect the slower progression of disease, allowing neurons to sequester tau into mature aggregates. In contrast, PSP progresses more rapidly^8,9^. Here, tau may accumulate as smaller, stress-associated aggregates, (potentially due to the availability of pS356 phosphorylation on the aggregates). Thus, differences in aggregate burden and morphology likely reflect not just aggregation propensity, but also disease stage, regional vulnerability, and cellular capacity to manage tau pathology.

An additional consideration is that differences in aggregate abundance may partly reflect differential neuronal survival. In Alzheimer’s disease, longer disease duration and slower neurodegeneration may allow neurons to persist despite tau pathology, providing the opportunity for aggregates to accumulate and mature. In contrast, in PSP, where disease progression is typically more rapid, early neuronal loss may limit the detectable aggregate burden at the time of sampling, rather than reduced aggregation per se. Further work with larger datasets could compare the tau aggregate signatures of fast *vs*. slow progressors within each tauopathy.

### Seeding competence of insoluble tau

While soluble aggregates are increasingly suspected to be the primary toxic species, functional assays have historically focused on insoluble material. To probe whether tau aggregates differ in their ability to template pathology, we measured the seeding capacity of sarkosyl-insoluble tau in 3R and 4R biosensor lines; AD and PiD strongly seeded 3R tau, CBD seeded 4R tau, and AD and PSP exhibited minimal 4R seeding. These findings suggest that seeding competence is not solely determined by isoform content or aggregate abundance but likely reflects underlying conformational or biochemical features of the aggregates themselves. The 3R cell line is much more sensitive than the 4R cell line^36^, which would explain why minimal seeding was detected with PSP and AD seeds. With CBD, on the other hand, tau may adopt a more seed-competent conformation, or be modified in a way that enhances templated aggregation.

### Implications for biomarker development and targeted therapeutics

This overall heterogeneity implies that tau aggregates play divergent roles across diseases depending on their molecular characteristics. These insights highlight the need for disease-specific approaches to both diagnosis and therapy. Soluble tau aggregates, when characterised by features such as phosphorylation state or morphology, may serve as more precise biomarkers than bulk tau levels. Likewise, tau-targeting interventions may benefit from tailoring to the specific aggregate species and mechanisms predominant in each tauopathy.

### Limitations of the study

While valuable for showing distinct morphologies of tau aggregates in disease-specific cases, this study is not free of limitations. While we focused on tau aggregates extracted from a single brain region, tau pathology is known to vary spatially, and regional differences in aggregate profiles may further inform disease mechanisms. The study design also does not capture how tau aggregation evolves over time. Moreover, the limited sample size (n = 4 for PiD and n = 5 cases for other disease groups) limits the generalisability of some findings and may reduce power to detect subtle differences across diseases or correlated within each disease group. Finally, while we identified correlations between tau aggregate features and inflammatory markers, causality cannot be inferred. Additional studies using longitudinal models or human-derived systems will be required to clarify the directionality and mechanisms underlying these associations.

In conclusion, we demonstrate that the small tau aggregates exhibit disease-specific signatures of their molecular and functional properties, which are associated with divergent cellular responses across tauopathies. By mapping these differences across structural, functional, and tissue-response domains, this work provides a framework for understanding tauopathy heterogeneity and identifying more precise, disease-informed biomarkers and therapeutic targets.

## RESOURCE AVAILABILITY

### Data and code availability

All original code has been deposited at Zenodo at DOI: 10.5281/zenodo.15635269 and is publicly available as of the date of publication. All data are available in the manuscript or the supplementary material. Raw data available on written request to DK.

## ACKNOWLEDGEMENTS

We thank the brain tissue donors and their families for their invaluable contribution to this study. This work is supported by the UK Dementia Research Institute through UK DRI Ltd, principally funded by the Medical Research Council. The Cambridge Brain Bank is supported by the NIHR Cambridge Biomedical Research Centre (NIHR203312). We thank the Cambridge Brain Bank and the Edinburgh Brain & Tissue bank for the provision of material.

DK holds a Royal Society funded Professorship. JBR is supported by the Wellcome Trust and NIHR Cambridge Biomedical Research Centre (NIHR203312). DC was supported by the Lady Edith Wolfson Junior Non-Clinical Research Fellowship awarded by the MND Association UK (Cox 971-799), and Discovery Early-Career Researcher Award from the Australian Research Council (DE240100707). WAM was supported by a Sir Henry Dale Fellowship jointly funded by the Wellcome Trust and the Royal Society (grant 206248/Z/17/Z). Further support was provided by the UK Dementia Research Institute (award number UK DRI-2010) through UK DRI Ltd, principally funded by the UK Medical Research Council. MH was supported by a Studentship funded by Aprinoia Therapeutics Limited.

For the purpose of open access, the authors have applied a CC BY public copyright licence to any Author Accepted Manuscript version arising from this submission.

## AUTHOR CONTRIBUTIONS

Conceptualization, D.B. and D.K; methodology, D.B., M.H., and D.C.; investigation, D.B., M.H., Y.W., E.F., J.Y.L.L. G.M. and S.B.; writing—original draft, D.B. and D.K.; writing—review & editing, D.B., M.H., Y.W., E.F., J.Y.L.L., G.M., J.B.R., A.Q., D.C., W.M., D.K.; funding acquisition, D.K.; resources, C.S.,

A.Q. and W.M.; supervision, W.M., and D.K.

## SUPPLEMENTAL INFORMATION

Document S1. Figures S1–S4

## METHODS

### Post-mortem brain tissue

Fresh-frozen post-mortem brain tissue samples from PiD and CBD donors were obtained from the Cambridge Brain Bank under the ethically approved protocol for ‘Neurodegeneration Research in Dementia’ (REC 16/WA/0240) and from AD, PSP and control donors from the Edinburgh Brain Bank with the approval from the East of Scotland Research Ethics Service (REC 21/ES/0087). The control group consisted of age-and gender-matched neurologically healthy controls with mild age-related pathology.

Written informed consent from the donors and/or their next of kin was provided as appropriate. Donor characteristics are summarised in Table 2.

### Preparation human brain tissue homogenate

Fresh-frozen brain tissue was homogenised using a method adapted from Goedert, et al.^46^, and as previously described^32^. Briefly, the tissue was homogenised at 4 °C in 10 volumes of filtered, ice-cold homogenisation buffer (10 mM Tris-HCl, 0.8 M NaCl, 1 mM EGTA, 0.1% Sarkosyl, 10% sucrose, 1x cOmplete^TM^ Ultra protease inhibitor mix and 1x PhosStop^TM^ phosphatase inhibitor mix; pH 7.32) using a VelociRuptor V2 Microtube Homogeniser (Scientific Laboratory Supplies) at 5 m/s, for two 20-s cycles with a 10-s rest in between. The homogenate was centrifuged (21,000 × g, 20 min, 4 °C), and the upper 90% of the supernatant was retained. The pellet was re-homogenised in 5 volumes of homogenisation buffer (5 m/s, 2× 20 s, 10 s rest), and again centrifuged (21,000 × g, 20 min, 4 °C). The upper 90% of this supernatant was removed, combined with the first supernatant, and aliquoted and frozen at -80 °C until used for further experiments. Total protein concentration was determined using a BCA assay (Thermo Fisher, Cat. No. 23227) as per the manufacturer’s instructions and was used for normalisation.

### Preparation of sarkosyl-insoluble tau from human brain tissue

Frozen brain tissue (120 mg) was homogenised in 10 volumes (v/w) of extraction buffer (800 mM NaCl, 10 mM Tris-HCl pH 7.4, 2.5 mM EDTA, 10% sucrose, 2% sarkosyl) supplemented with Pierce Protease and Phosphatase Inhibitor Mini Tablets (Thermo Scientific, A32959). Homogenisation was performed using a T 10 basic ULTRA-TURRAX (IKA 9993737002). The resulting homogenate was incubated at room temperature for 1 h on a roller before centrifugation at 13,000 g for 20 minutes at room temperature. The supernatant was filtered through a 0.45 μm cell strainer and subjected to ultracentrifugation (124,500x g, 1 h at room temperature) using a TLA-55 rotor in a Beckman Coulter Optima MAX-XP ultracentrifuge. The resulting pellets were pooled, resuspended in extraction buffer, and re-centrifuged under the same conditions. The final pellet was resuspended in 50 mM Tris-HCl, 150 mM NaCl, followed by a further ultracentrifugation step at (124,500x g, 1 h). The final pellet was resuspended in 250 μL buffer per gram of starting tissue weight and sonicated in a water-bath sonicator for 20 cycles of 15-s on / 5-s off. The relative concentration of sarkosyl-insoluble tau was determined using a Human Tau ELISA Kit (abcam, ab273617).

### Antibodies

All antibodies were obtained as a carrier-protein-free composition. All tau antibodies were monoclonal to enable specific aggregate detection. To ensure the detection of aggregates, the same antibody was used for capture and detection in Simoa and SiMPull assays except for p53 where DO-1 was used for capture and PAb240 was used for detection. The antibodies used in the study are listed in Table 3.

**Table 3:** Antibodies used in the study. Mono: monoclonal, H: human, M: mouse, B: bovine, R: rat, Co-loc: co-localisation SiMPull experiments, SR: super-resolution experiments.

| Type | Antibody | Manufacturer | Cat. No. | Clonality | Reactivity | Simoa | Co-loc | SR |
| --- | --- | --- | --- | --- | --- | --- | --- | --- |
| Tau | HT7 | Invitrogen | MN1020(b) | mono | H | x | x | x |
|  | Tau13 | Santa Cruz | sc-21796 | mono | H | x | x |  |
|  | Tau46 | Sigma | T9459 | mono | H, M, B, R | x | x |  |
|  | 4R | Cell Signaling | 57664SF | mono | H, M, R | x |  |  |
|  | AT8 | Invitrogen | MN1020(b) | mono | H, M | x | x | x |
|  | T181 | abcam | ab236458 | mono | H | x |  |  |
|  | T217 | abcam | ab314558 | mono | H | x |  |  |
|  | S356 | abcam | ab196367 | mono | H | x |  |  |
| Other | AL177 | AdipoGen | AL177 | poly | H, M |  |  | x |
|  | 6E10 | BioLegend | 803001 | mono | H | x |  |  |
|  | Ubiquitin | abcam | ab230145 | mono | H, M |  | x |  |
|  | DO-1 | Abcam | ab1101 | mono | H | x |  |  |
|  | PAb240 | Novus Bio. | NB200-103B | mono | H | x |  |  |

### Labelling of antibodies

#### Fluorophore

Carrier protein-free antibodies were fluorescently labelled with AF647 or AF488 using the Alexa Fluor Conjugation Kit (Fast) - Lightning-Link (abcam, ab269823 and ab236553) according to manufacturer’s instructions. Excess fluorophore was removed using a 40 kDa Zeba Spin desalting column (Thermo Fisher, A57756) followed by a 50 kDa Amicon Ultra Spin column (Merck, UFC5050). The dye-labelled antibodies were stored at 4 °C. The protein concentration and degree of labelling were determined through absorption at 280 nm and 494 nm (AF 488) or 650 nm (AF647), taking into account the absorption of the dyes at 280 nm. The degree of labelling was within the manufacturer’s recommendation of 3-9 moles of dye per mole of antibody.

#### Biotin

Antibodies were biotinylated using the Biotin Conjugation Kit (Fast, Type B) - Lightning-Link (abcam, ab201796) following manufacturer’s instructions. Excess biotin was removed using a 40 kDa Zeba Spin desalting column (Thermo Fisher, A57756) followed by a 50 kDa Amicon Ultra Spin column (Merck, UFC5050). The biotin-conjugated antibodies were stored at 4 °C. The protein concentration was determined through absorption at 280 nm.

#### DNA

For DNA-PAINT experiments the DBCO-conjugated docking strand DBCO-concatDS2 (DBCO TEG-AAACCACCACCACCACCACCACCACCACCACCACCA) and atto655 conjugated imaging strand (TGGTGGT-AminoC7-atto655) were purchased from ATDBio (Southampton, UK). DBCO-concatDS2 was synthesised on the 1.0 μmol scale and purified by HPLC. The dye-labelled imaging strand was synthesised on the 0.2 μmol scale and purified by double HPLC. The lyophilised oligonucleotides were dissolved in 18.2-MΩ·cm water (filtered by 0.02-μm filter (VWR, 516-1501)) the concentration determined by A260, and aliquoted and stored at −20 °C. The detection antibody was labelled with the DNA docking strand (DBCO-concatDS2, ATDBio). The antibody was prepared using the SiteClick Antibody Azido Modification kit (Invitrogen, S20026) following manufacturer’s instructions. To the azido-modified antibody, 10 molar equivalents of docking strand were added and incubated overnight at 37 °C. Excess docking strand was removed using an Amicon spin filter (100 kDa MWCO). The concentration of antibody and degree of labelling (typically 3-4 docking strands per antibody) was determined by A280 and A260/A280.

### Single-molecule imaging

#### Single-Molecule Pull Down

Coverslips were passivated using polyethylene glycols (PEGs) and biotin-conjugated PEGs as previously described^32^. Diffraction-limited and two-colour co-localisation experiments were performed as previously described^32^ with small modifications. Briefly, the volume typically used per well was 10 µL, unless otherwise indicated. Each washing cycle consisted of two washes with PBST (0.05% Tween in PBS) followed by one wash with PBST containing 1% Tween which was incubated for 5 minutes. Throughout the assay the coverslip was kept in a humidity chamber. First, NeutrAvidin (Thermo Fisher) was added to each well at 0.2 mg/mL in PBST and incubated for 10 min. After a washing cycle, 10 nM biotinylated antibody in 1 mg/mL BSA in PBS was incubated for 10 minutes, followed by another washing cycle, and a 10 min blocking step (1 mg/mL BSA in PBS). Next, 10 µL of sample (diluted in PBS, for tau 1:10, for 6E10 1:5, for ASC: 1:20) was added to each well and left to incubate for 1h at room temperature. Subsequently, another washing cycle was performed and then fluorescently labelled antibody (AT8: 4 nM, HT7: 2 nM, 6E10: 1 nM, ASC: 5 nM, all in 0.1 mg/mL BSA in PBS) was added and incubated for 15 minutes in the dark. A last washing cycle was performed. Lastly, 10 µL of PBS was added to each well and the wells were sealed with a clean glass coverslip.

For co-localisation experiments two fluorescently labelled antibodies (AF647 and AF488) were mixed at 2 nM each and incubated at the same time. All other steps were performed as described above.

For DNA PAINT experiments, DNA-labelled detection antibody (DBCO TEG-AAACCACCACCACCACCACCACCACCACCACCACCA, ATDBio) was used and the corresponding imaging strand (TGGTGGT – Amino C7 – Atto655, ATDBio) was added at the last step (3 µL, 1 nM in PBS).

#### Microscope setup

Imaging was performed on a home-built total internal reflection fluorescence (TIRF) microscope, consisting of an inverted Ti-2 Eclipse microscope body (Nikon) fitted with a 1.49 N.A., 60x TIRF objective (Apo TIRF, Nikon) and a perfect focus system. The coverslip was illuminated using a 638 nm laser (Cobolt 06-MLD-638, HÜBNER) and a 488 nm laser 488 nm (Cobolt 06-MLD-488, HÜBNER).

#### Diffraction-limited image acquisition and analysis

For each well, 16 field-of-views (FOVs) were typically imaged, each comprising 50 frames of 50 ms exposure. To avoid any bias, an automated script (*Micro-Manager*^47^) was used to collect images in a 4x4 grid. Individual fluorescent spots were quantified using a python-based adaptation of *ComDet*^48^. Briefly, to minimise background fluctuations a mean intensity projection was prepared from the last 40 frames of each FOV. Thresholds were then optimised by comparing the number of particles identified in positive and negative control images, such that the threshold was set to the lowest value at which the negative control had less than 1% of the positive control. This threshold was then used along with a particle size estimate of 4 to identify fluorescent spots. The mean intensity of each spot was measured using *scikit-image*.

For co-localisation experiments, images were acquired sequentially in each excitation channel in a 4x4 grid. Spot detection was performed independently on both channels using *ComDet*, generating separate lists of centroid coordinates for each channel. Co-localised spots were then identified based on a Euclidean distance threshold of 4 pixels, with each spot in one channel assigned to the nearest spot in the opposing channel if it fell within this radius. In cases where multiple spots met this criterion, the closest spot was selected as the co-localised pair. To estimate chance co-localisation, the centroid coordinates of the second channel were transposed along the x-dimension, and the distance-based analysis was repeated. The proportion of co-localised spots was then calculated as the number of matched spots divided by the total number of detected spots in the given channel.

#### Super-resolution image acquisition and analysis

For DNA-PAINT imaging, SiMPull coverslips were prepared as described but with modifications. After the sample incubation, the coverslip was washed two times with PBST and one time with 1% Tween in PBS, followed by a 30 min blocking step with 1 mg/mL BSA in PBS. The blocking solution was then removed, and DNA-labelled detection antibody was incubated for 15 min (3 nM in 0.1 mg/mL BSA in PBS). The coverslip was again washed two times with PBST, one time with 1% Tween in PBS and one time with PBS before the corresponding imaging strand (TGGTGGT-AminoC7-atto655, ATDBio) was added at the last step (5 µL, 1 nM in PBS). The coverslip was sealed with a clean coverslip. Imaging was performed using a 638 nm laser at 190 mW. Images of typically 4 FOVs were collected in a 2-by-2 grid using an automated script (*Micro-Manager*) to avoid any bias in the selection of FOVs. Images were acquired for 8000 frames of 100 ms exposure.

Super-resolution images were reconstructed using the *Picasso* package^49^. Briefly, after discarding the first 100 frames, localisations were identified, fitted, and then corrected for microscope drift using redundant cross-correlation. Localisations were then filtered for precision <30 nm. To obtain aggregate clusters, *DBSCAN* was applied using the *scitkit-learn* package^50^ (radius of 0.3 (∼35nm) and minimum density of 5). Each aggregate was then measured using a combination of scikit-image^51^ for basic region properties such as perimeter, area, and eccentricity, and *SKAN*^52^ for skeletonised length whereby the length of each aggregate is reported as the summed branch distance. Finally, super-resolved images were rendered using the inbuilt *Picasso*^49^ functionality.

### Simoa

#### Simoa bead conjugation

For the conjugation of antibody to the Simoa beads the manufacturer’s instructions were followed, using a two-step EDC protocol that couples the antibody primary amino groups to the activated carboxyl groups on the beads. The reaction was performed using 0.3 mL of bead suspension with 0.2 mg/mL antibody and 0.3 mg/mL EDC at 4 °C. Briefly, 100 μg of antibody was buffer exchanged to the bead conjugation buffer (Quanterix) and the concentration was adjusted to 0.2 mg/mL. The beads were washed and activated with EDC on a rotator for 30 min at 4 °C. Subsequently, antibody was added to the beads and the mixture was incubated on a rotator for 2h at 4 °C. Excess antibody was removed by washing the beads before the beads were blocked for 45 min. To determine the antibody coating efficiency, the absorbance at 280 nm was measured in the supernatant and washes. A coupling efficiency of 90% was achieved. Beads were stored as a pellet at 4 °C.

#### Simoa assay

The Simoa assays were run on a 96-well plate as a 3-step assay as previously described^33^. Briefly, 100 μL of sample diluted in Quanterix Tau 2.0 sample diluent was incubated with 25 μL of antibody-conjugated paramagnetic beads (30 minutes, 800 rpm, 30 °C). After an automated washing cycle, 100 μL biotinylated detector antibody (0.3 ng/mL in Quanterix Detector Diluent) was added to each well and incubated (10 minutes, 800 rpm, 30 °C). After a further automated washing cycle, 100 μL SBG (150 pM in Quanterix SBG Diluent) was added to each well and incubated (10 minutes, 800 rpm, 30 °C). The plate was then loaded onto the Simoa SRX instrument (Quanterix) and new RGP substrate, which has been incubated at 30 °C for 2h, was added.

### Cellular markers

ELISAs were used to determine the levels of total tau (Human Tau ELISA Kit, abcam, ab273617), 4R tau (Human Tau (4R isoforms) ELISA Kit, abcam, ab309282), MLKL (Human MLKL ELISA Kit, antibodies.com, A82371), BAX (Human BAX SimpleStep ELISA Kit, abcam, ab199080), Iba1 (Human Iba1 ELISA Kit, antibodies.com, A5204), and GFAP (Human GFAP ELISA Kit, abcam, ab223867). All ELISA assays were performed following manufacturer’s instructions. The assays were read using absorbance at 450 nm and concentrations were determined by fitting a 4PL curve to the standards. ASC speck levels were determined through DNA-PAINT, following the SiMPull protocol described, which was adapted from Fertan *et al*.^40^. Levels of p53 were determined using a Simoa assay described in Wu *et. al.*^39^.

### Full-length, wild-type 3R and 4R tau HEK293 seeding assay

The full-length wild-type tau seeding assay was performed as previously described^36^. Briefly, HEK293 cells expressing either eGFP-0N3R tau or eGFP-0N4R tau were used to assess 3R/4R specific seeding and seeded with sonicated (20 cycles of 15-s on/5-s off) sarkosyl insoluble tau at 13.75 ng/mL. Black, glass-bottom 96-well plates (Ibidi) were first coated with Poly-D-Lysine (Gibco, A3890401) for 30 minutes at 37 °C, followed by three washes with 100 μL per well of PBS. Tau assemblies were prepared at twice the final desired concentration in OptiMEM Reduced Serum Media (Gibco, 31985062) and mixed with Lipofectamine 2000 (Invitrogen, 11668019) at a 1:50 dilution in a total volume of 50 μL per well. This mixture was incubated for 20 minutes at room temperature. Meanwhile, cells were trypsinised and counted using a Countess II Automated Cell Counter (Invitrogen), then diluted to 40,000 cells/mL in OptiMEM. For each well, 50 μL of the cell suspension was added to 50 μL of the tau-Lipofectamine mixture, yielding final concentrations of 1X tau, 1:100 Lipofectamine 2000, and 20,000 cells per well. Plates were incubated for 1 hour, after which 100 μL per well of DMEM supplemented with 10% FBS was added. Cells were incubated for 72 hours to allow seeding before imaging on a Ti2 High Content Microscope (Nikon) with a 10x lens and analysis using the NISElements software.

### Production of 3R and 4R tau

Purified 3R and 4R tau aggregates and monomers for the validation of the 4R tau aggregate Simoa assay were produced as previously described^36^. Human 6xHis-tagged 0N3R and 0N4R tau constructs in pRK172 were expressed in *E. coli* BL21 (DE3) and grown (37°C, 200 rpm) in 2XTY media to OD600 = 0.5 before induction with 1 mM IPTG and overnight incubation at 16 °C, 200 rpm. Cells were harvested, lysed in buffer (25 mM HEPES pH7.4, 300 mM NaCl, 20 mM Imidazole, 1 mM Benzmidine, 1 mM PMSF, 14 mM β-mercaptoethanol, 1% NP-40 with Pierce protease and phosphatase inhibitor tablets (Thermo Scientific, A32959), and lysates cleared by ultracentrifugation. Tau was purified using His-tag affinity chromatography (His/TRAP HP column, GE Healthcare) and a Nickel affinity column, followed by size-exclusion chromatography on a Superdex 200pg column equilibrated in TBS + 1 mM DTT. Tau aggregation was induced by incubating 60 μM recombinant tau with 20 μM heparin in aggregation buffer at 37 °C, shaking at 700 RPM for 72 h. Aggregation was monitored using 15 μM Thioflavin T fluorescence.

### Western blots

Sarkosyl-insoluble tau, from the equivalent of 4mg of brain tissue, was diluted in TBS and boiled for 5 minutes at 95°C in 1X NuPage LDS Sample Buffer (ThermoFisher) + 4% BME. Samples were run on 4-20% Novex Tris-Glycine gels (ThermoFisher Scientific) for 50 minutes at 200 V and transferred onto PVDF membranes using the Trans-Blot Turbo Transfer system (BioRad). Membranes were blocked in TBS + 0.1% Tween-20 + 5% milk for 1 hour and incubated overnight at 4°C in primary antibody. Following washes in TBS + 0.1% Tween-20, blot was incubated in anti-mouse StarbrightBlue 700 (BioRad #12004158, 1:2500) or anti-rabbit DyLight 800-conjugated secondary antibody (Cell Signaling Technology, 5151, 1:10,0000) for 45 minutes. Signal was visualized using near IR fluorescence detection (ChemiDoc MP, BioRad). The following primary antibodies were used for immunoblotting: Tau 12 (Sigma-Aldrich, MAB2241, 1:2000); BR134 (in-house, rabbit polyclonal, 1:4000); RD3 (Sigma-Aldrich, 05-803, 1:1000); and Tau 4R (Cell Signaling Technology; 30328).

### Data analysis

#### Radar plot of aggregate features

To visualise disease-specific molecular profiles, we generated a radar plot comparing 15 aggregate-related features across tauopathies (small aggregates in brain homogenate). All values were min–max normalised to a [0,1] range using *MinMaxScaler* from *scikit-learn*. For each disease group, we computed the mean of each feature using the scaled data.

#### Linear discriminant analysis

To identify disease-specific patterns of tau aggregate features, we performed linear discriminant analysis (LDA) using *scikit-learn*. The analysis included 13 brain-homogenate aggregate-associated features (mean length of HT7 aggregates, percentage of long aggregates, localisation density of long aggregates, levels of tau aggregates in brain homogenate (HT7 Simoa), levels of total tau in brain homogenate, percentage of 4R aggregates in brain homogenate, levels of AT8, pT181, pT217 and pS356 (Simoa), percentage of aggregates with Tau46 epitope, percentage of aggregates with Tau13 epitope, and percentage of AT8 aggregates positive for ubiquitin. Data points were labelled according to disease group and LDA was used to project the data onto a lower-dimensional space that maximally separates the disease classes based on between-group variance. Disease group separation and feature contributions were further quantified using the explained variance ratio and loading coefficients for each discriminant axis. The top contributing features to each linear discriminant were identified based on absolute loading weights. The first two discriminant axes were visualised in a 2D scatter plot with group-wise colour coding.

#### Relative change matrix for stress and inflammatory markers

To compare disease-specific changes in stress and inflammatory markers, we computed the relative level of each marker in tauopathy groups versus healthy controls (HC). Marker values were first normalised to total protein concentration. The relative changes were scaled to the maximum absolute change observed across all groups, resulting in a normalised range of –1 to +1 per marker.

#### Partial correlation analysis

To investigate associations between tau aggregate features and cellular stress or inflammatory markers while accounting for disease group differences, we computed pairwise partial correlation coefficients. For each variable pair, we used ordinary least squares regression to control for diagnosis (encoded as dummy variables), and calculated Pearson’s correlation between the residuals. This yielded a partial correlation matrix. Only correlations passing a significance threshold (p < 0.05) were annotated in the resulting heatmap.

### Statistical analysis

Details for the statistical analysis for each experiment can be found in the respective figures and figure legends, including the statistical tests used, sample size and definition of significance. Statistical analysis was performed using Python (*scikit-learn*) and GraphPad Prism 10 (version 10.4.1). For all analyses involving the comparison of one variable, one-way ANOVAs were used followed by Tukey’s multiple comparisons post hoc corrections. For analysis of tau seeding activity in biosensor assays, a nested one-way ANOVA was applied. For clarity, only statistically significant differences are shown in the figures. A level of *p* < 0.05 was used to designate significant differences.

---

**Supplementary Figure 1.**
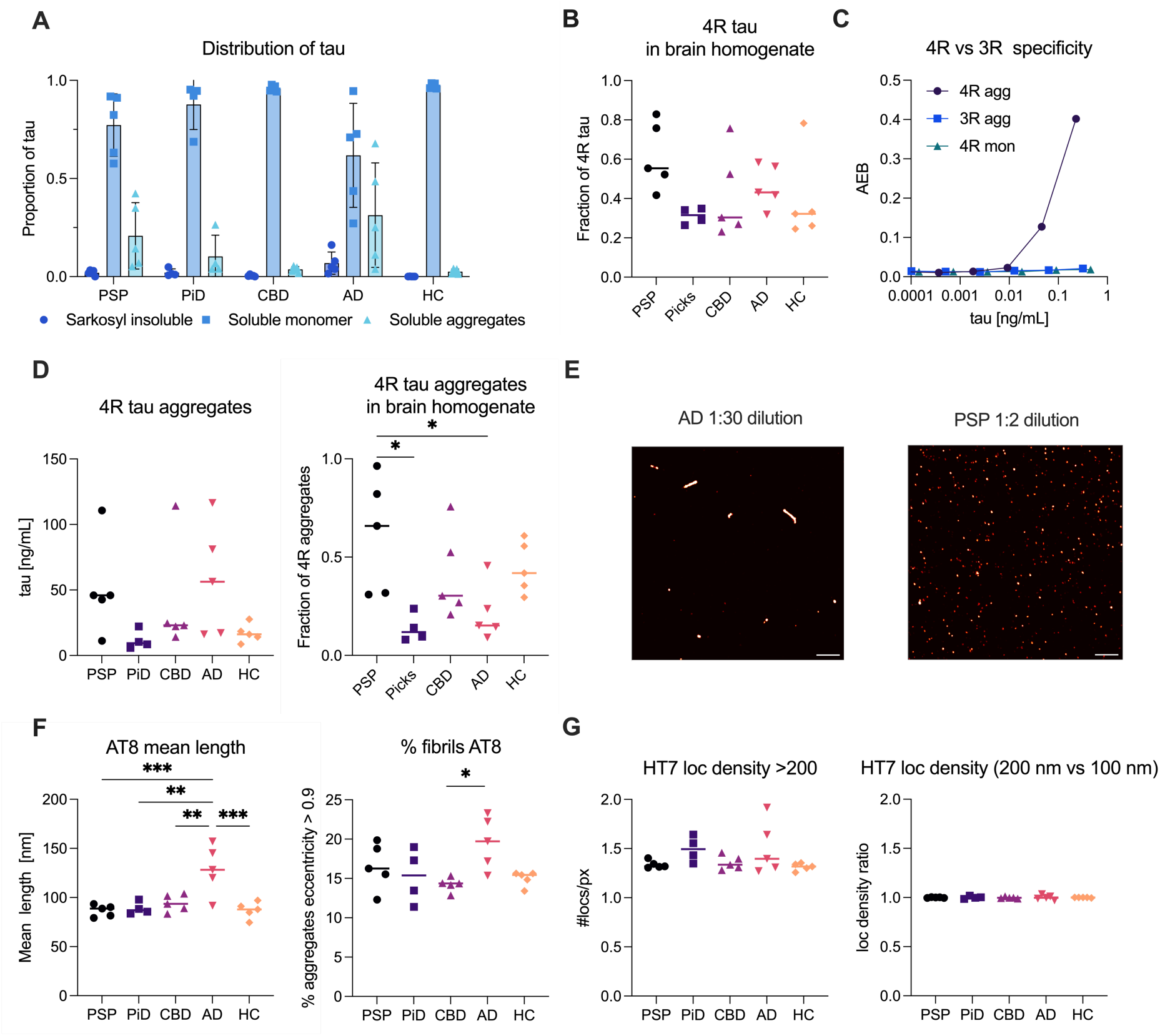
: Quantitative and morphological characterisation of small tau aggregates in brain homogenate. (A) Proportion of tau present either as monomer, small aggregates or sarkosyl insoluble tau by patient. Each point represents one case (PiD: n = 4 per group, rest n = 5 per group) and the mean ± S.D. is shown. (B) Fraction of 4R tau in homogenate (levels of 4R tau normalised to total tau levels in homogenate fraction) of the individual patients. (C) Validation of the Simoa-based 4R tau aggregate assay, showing high specificity and concentration-dependency for 4R tau aggregates but not for 4R tau monomers or 3R tau aggregates. (D) Levels of small 4R tau aggregates in the brain homogenate fraction. (E) Representative DNA-PAINT super-resolved image of an AD sample at 1:30 dilution (compared to the normal 1:10) showing the presence of long aggregates and of a PSP sample at 1:2 dilution, showing the presence of small, round aggregates in a crowded image. Scale bar = 1 µm. (F) Mean length of small AT8-positive aggregates and percentage of fibrillar AT8-positive aggerates (eccentricity > 0.9). (G) Localisation density of tau aggregates in an HT7 DNA-PAINT experiment. Each point represents one case (PiD: n = 4 per group, rest n = 5 per group).

**Supplementary Figure 2.**
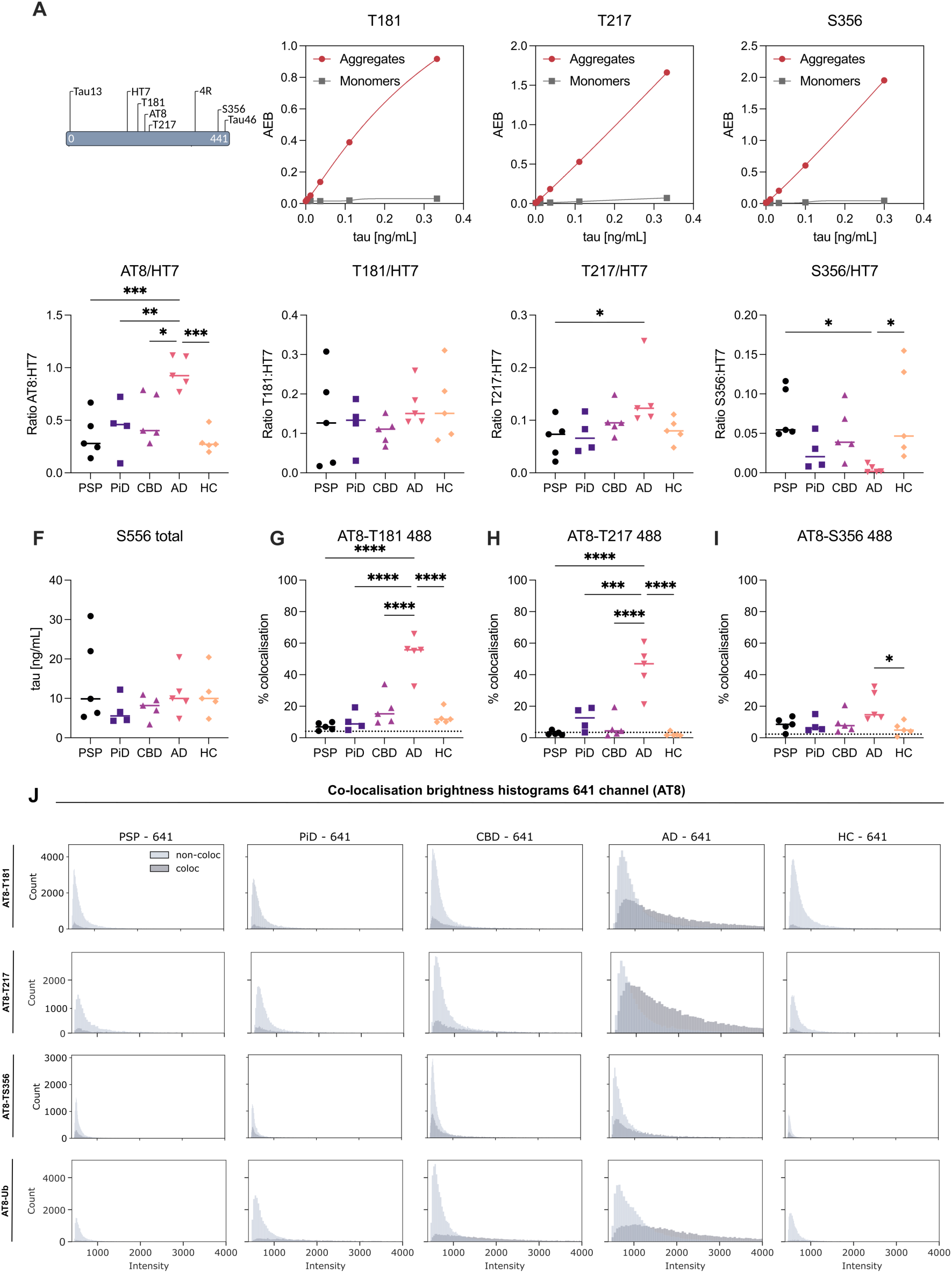
: **Phospho-epitope profiling and co-localisation of small tau aggregates.** (A) Validation of Simoa assays for the detection of tau aggregates phosphorylated at T181, T217 and S356. (B–E) Aggregate-specific Simoa assays quantifying the proportion of tau aggregates positive for phospho-epitopes AT8 (pS202/pT205), pT181, pT217, and pS356, expressed as ratios over HT7 aggregates (pan tau). (F) Total levels of S356 phosphorylation (monomeric and aggregated tau) measured by Simoa. (G–I) Percentage of co-localisation of pT181, pT217, or pS356 (488 channel) with AT8-positive tau aggregates as determined by two-colour single-molecule co-localisation microscopy. (J) Histograms of fluorescence intensity (AT8-641 channel) of AT8-positive aggregates, stratified by co-localisation with secondary phospho-epitopes (488 channel). Each row corresponds to a co-labelling condition (AT8–T181, T217, S356, Ubiquitin), and each column to a disease group (PSP, PiD, CBD, AD, HC). Co-labelled and non-co-labelled populations are shown in dark and light grey, respectively. Each point represents one case (n = 4 for PiD, n = 5 for other disease groups). For (B – I) one-way ANOVA with Tukey’s post hoc test; \**p* < 0.05, \*\**p* < 0.01, \*\*\**p* < 0.001.

**Supplementary Figure 3.**
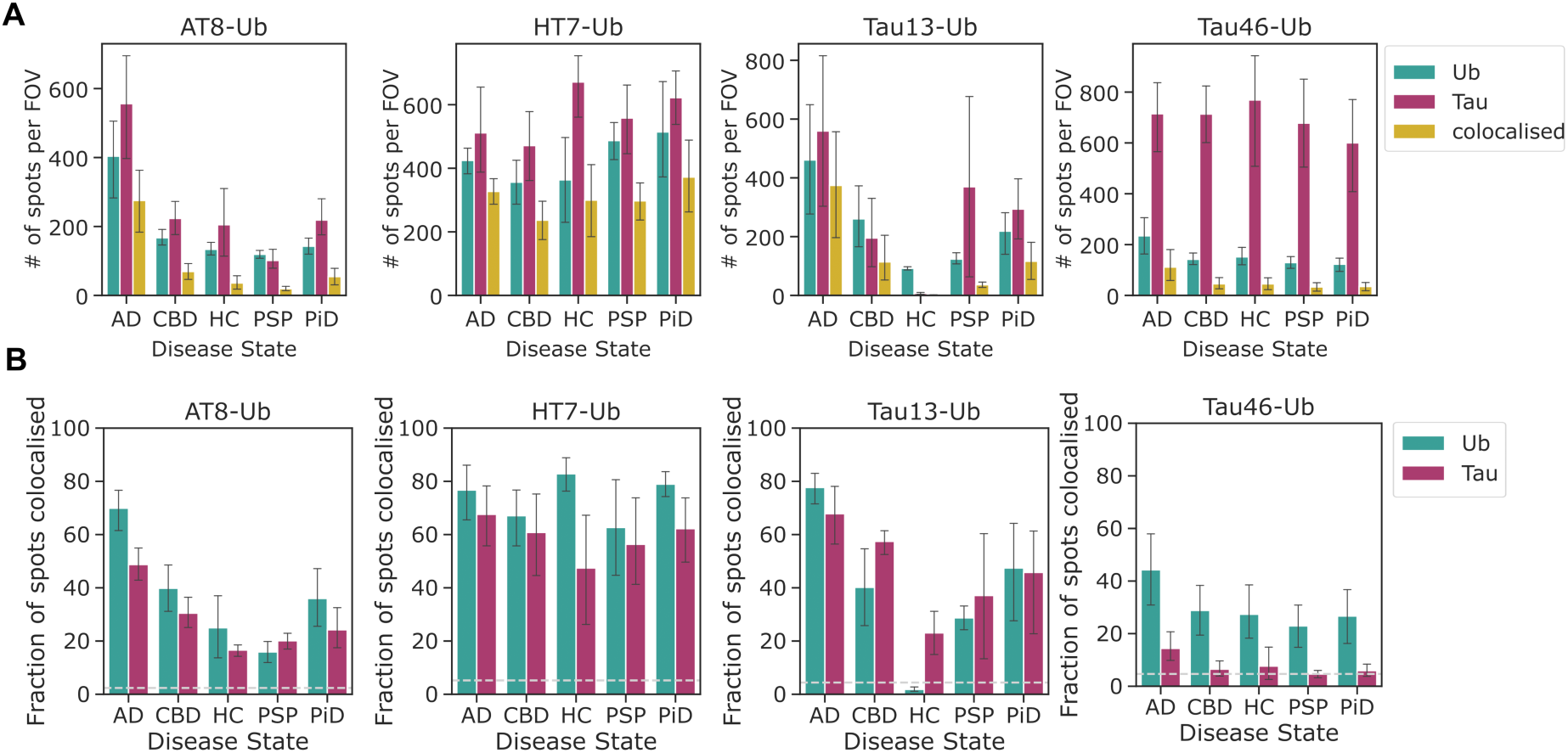
: C**o-localisation of tau and ubiquitin in brain homogenate across tauopathies.** (A) Number of spots per field-of-view (FOV) detected by SiMPull using tau (AT8, HT7, Tau13, Tau46 in the 641 channel) and ubiquitin antibodies (488 channel), with spatially overlapping signals indicating co-localised aggregates. (B) Fraction of tau or ubiquitin spots co-localised with the respective partner, plotted separately for each antibody pair. Dotted lines denote expected random co-localisation rates. Each bar represents mean ± SD (n = 4 for PiD, n = 5 for other disease groups).

**Supplementary Figure 4.**
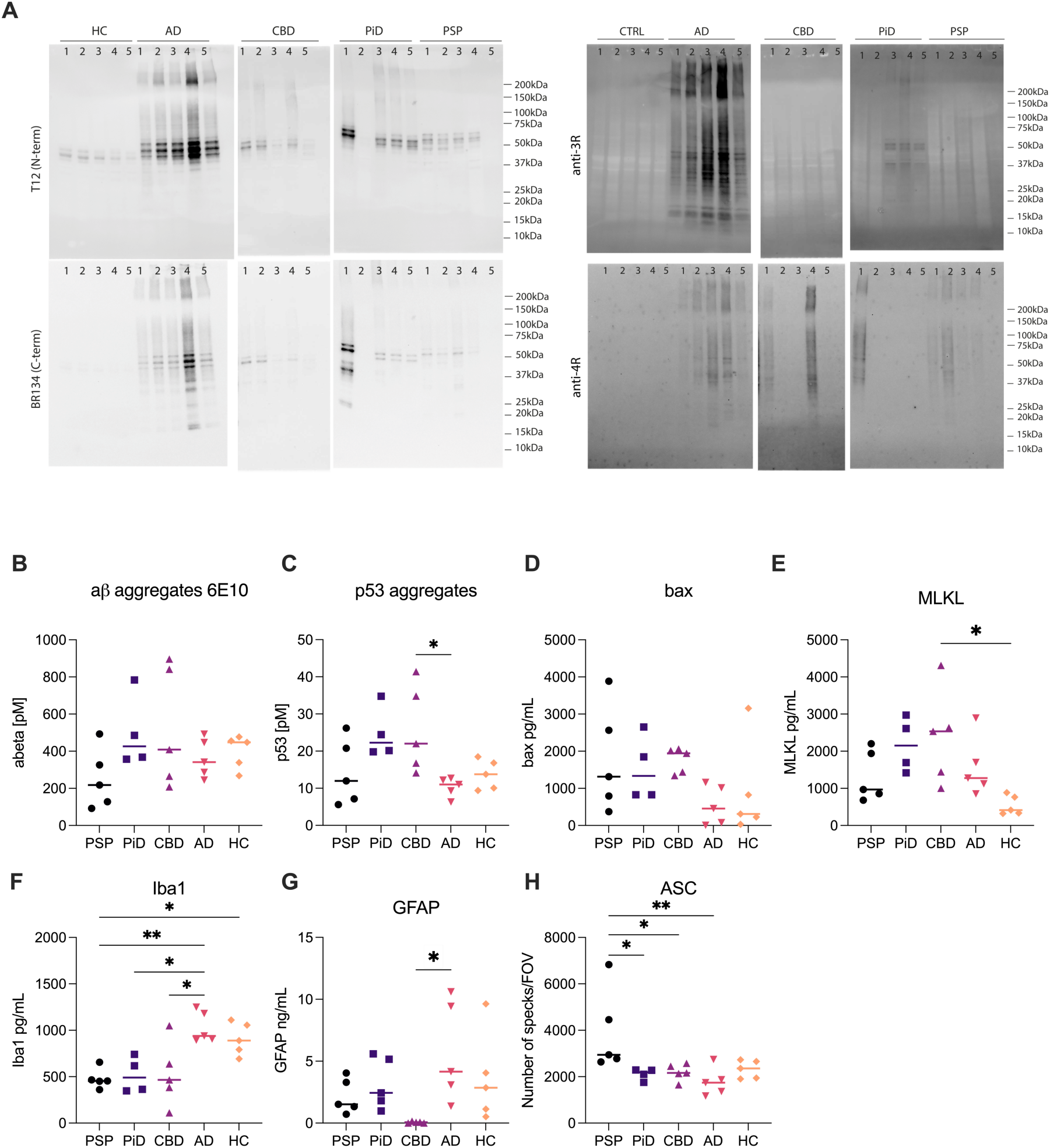
: Disease-specific profiles of stress and inflammatory markers across tauopathies. (A) Western blot analysis of sarkosyl insoluble fraction. (B–H) Quantification of cell stress and inflammatory markers in brain homogenate from AD, PSP, CBD, PiD, and HC. Markers include: (B) Aß aggregates (Simoa), (C) p53 aggregates (Simoa), (D) Bax (apoptosis, ELISA), (E) MLKL (necroptosis, ELISA), (F) Iba1 (microglial activation, ELISA), (G) GFAP (astrocytic activation, ELISA), and (H) ASC specks (inflammasome activation, SiMPull). Each point represents an individual donor (n = 4 for PiD, n = 5 for other disease groups); horizontal bars indicate group medians. One-way ANOVA with Tukey’s post hoc test; \**p* < 0.05, \*\**p* < 0.01.

